# Parkinson’s disease-linked D620N mutation selectively alters the brain-specific protein interactome of VPS35

**DOI:** 10.64898/2026.04.09.717005

**Authors:** Erin T. Williams, Xi Chen, Jordan Rowlands, Md. Shariful Islam, Maxwell Frye, Darren J. Moore

## Abstract

Mutations in several genes are known to cause familial forms of Parkinson’s disease (PD), including mutations in the *vacuolar protein sorting 35 ortholog* (*VPS35*) gene linked to late-onset, autosomal dominant PD. *VPS35* encodes a core subunit of the retromer complex which functions in endosomal sorting and recycling. It remains unclear how the pathogenic D620N mutation in VPS35 disrupts retromer function to induce neurodegeneration in PD. Using cell-and rodent-based models expressing D620N VPS35, we performed interactome proteomics to identify alterations underlying the pathogenic effects of D620N VPS35 in PD. Using overexpression of VPS35 variants in HEK-293T cells, we conducted tandem affinity purification (TAP) or co-immunoprecipitation (co-IP) with protein chemical crosslinking to determine the native and non-native protein interactomes of wild-type (WT) and D620N VPS35, respectively. Notably, we can confirm the reduced interaction of D620N VPS35 with components of the WASH complex. Additionally, using a viral-mediated gene transfer model of human D620N VPS35 overexpression in adult rat brain, we identify the first brain-specific protein interactome of VPS35. These overexpression models reveal remarkably similar interaction profiles of WT and D620N VPS35, suggesting that the D620N mutation has a subtle effect on the overall VPS35 protein interactome. We also conducted proteomic analysis of brain tissue from a *D620N VPS35* knockin (KI) mouse model that expresses VPS35 at endogenous levels. Using co-IP from hemi-brain or striatal extracts of *WT* and *D620N VPS35* KI mice, we reveal a high degree of similarity between the brain interactomes of WT and D620N VPS35, further suggesting a subtle effect of the D620N mutation on VPS35 protein interactions. Notably, in both hemi-brain and striatum, we find a selective decrease in the interaction of two known interactors, TBC1D5 and VPS29, with D620N VPS35. We also performed global proteomic analysis of striatal tissue from *D620N VPS35* KI mice and reveal a high degree of similarity between WT and D620N, further suggesting a subtle effect of this mutation. Together, our study provides a comprehensive evaluation of the VPS35 protein interactome and reveals a selective effect of the PD-linked D620N mutation in mammalian cells and brain. Our study provides key insight into the mechanisms of retromer dysfunction in *VPS35*-linked PD.

## Introduction

Parkinson’s disease (PD) is the most common neurodegenerative movement disorder that is neuropathologically characterized by the progressive loss of dopaminergic neurons in the substantia nigra pars compacta, leading to a reduction of striatal dopamine, and the accumulation of protein inclusions, termed Lewy bodies, containing misfolded α-synuclein, within the midbrain (1,2). The loss of dopamine in the striatum leads to the development of cardinal motor symptoms including tremor, rigidity, bradykinesia and postural instability (1,2). Although the majority of PD cases are considered sporadic with no clear cause, 5-10% of cases are hereditary (3). To date, mutations in more than 20 genes have been shown to cause familial forms of PD, including autosomal dominant mutations in *VPS35* (4–6).

Several missense mutations in *VPS35* have been associated with the development of PD worldwide, however the Asp620Asn (D620N) mutation has become the focus of several studies due to its clear segregation with disease in families with PD (7,8). PD subjects harboring heterozygous mutations in *VPS35* are clinically indistinguishable from sporadic patients, however neuropathology associated with *VPS35* mutations remain unclear due to limited postmortem neuropathology analysis (5,6,9). The mechanisms through which the D620N mutation exerts its pathogenic effects in PD remain unclear. Typically, autosomal dominant mutations cause pathogenicity through a toxic gain-of-function mechanism, however, they may also act through partial loss-of-function or dominant-negative mechanisms. It remains unclear through which of these mechanisms D620N VPS35 imparts pathogenic effects, however, multiple studies suggest that D620N VPS35 may induce defects in the autophagy/lysosomal pathway, mitochondrial function, Wnt/β-catenin signaling, and/or neurotransmission, ultimately leading to neurodegeneration (8,10,11). Interestingly, the molecular mechanisms underlying the dysfunction of these pathways have yet to be elucidated.

To that end, we hypothesized that the D620N mutation in VPS35 disrupts protein-protein interactions, which impairs retromer function and causes neurodegeneration. Using overexpression of VPS35 in HEK-293T cells, we conducted tandem affinity purification (TAP) and co-immunoprecipitation (co-IP) to determine native and non-native protein interactions of wild-type (WT) and D620N VPS35, respectively. Additionally, using a viral-based model of human D620N VPS35 overexpression in adult rats, we identify brain-specific interacting proteins of VPS35 variants. We also conducted VPS35 interactome analysis of brain tissue from a *D620N VPS35* knockin (KI) mouse model that expresses VPS35 at endogenous levels. Notably, in both hemi-brain and striatum, we identify a decrease in the interaction of TBC1D5 and VPS29 with D620N VPS35 compared to WT VPS35. Additionally, we performed global proteomic analysis on striatal tissue of *VPS35* KI mice to determine if the D620N mutation alters proteins levels in the brain. Taken together, our different models show remarkably similar interaction profiles of WT and D620N VPS35, suggesting that the D620N mutation has a subtle effect on VPS35 protein interactions and global protein levels. Interestingly, we identify a novel interaction deficit between D620N VPS35 and two well-defined binding partners of the retromer, which could contribute to retromer disfunction and VPS35-induced neurodegeneration.

## Results

### Optimization of tandem affinity purification methodology for VPS35 and the retromer

To identify native protein complexes of VPS35 variants, we employed the use of tandem-affinity purification (TAP) methodology. A TAP tag consisting of distinct streptavidin-binding and calmodulin-binding peptides were fused in frame to the C-terminus of VPS35. TAP has several advantages, including detergent-free pulldown, allowing for the preservation of native protein complexes, and two consecutive purification steps based on high affinity for streptavidin and calmodulin, which serve to decrease background contaminants (12,13). Using Agilent’s InterPlay Mammalian TAP System kit, TAP-tagged WT VPS35 was first tested for expression in both HEK-293T and SH-SY5Y cells (**Figure 1A**). TAP-tagged VPS35 produces an expected protein at ∼100 kDa detected by an anti-TAP tag antibody, which migrates slightly above the endogenous VPS35 protein but at modestly lower steady-state levels. HEK-293T cells were used for all subsequent TAP experiments owing to their higher transfection efficiency and expression levels.

**Figure 1.**
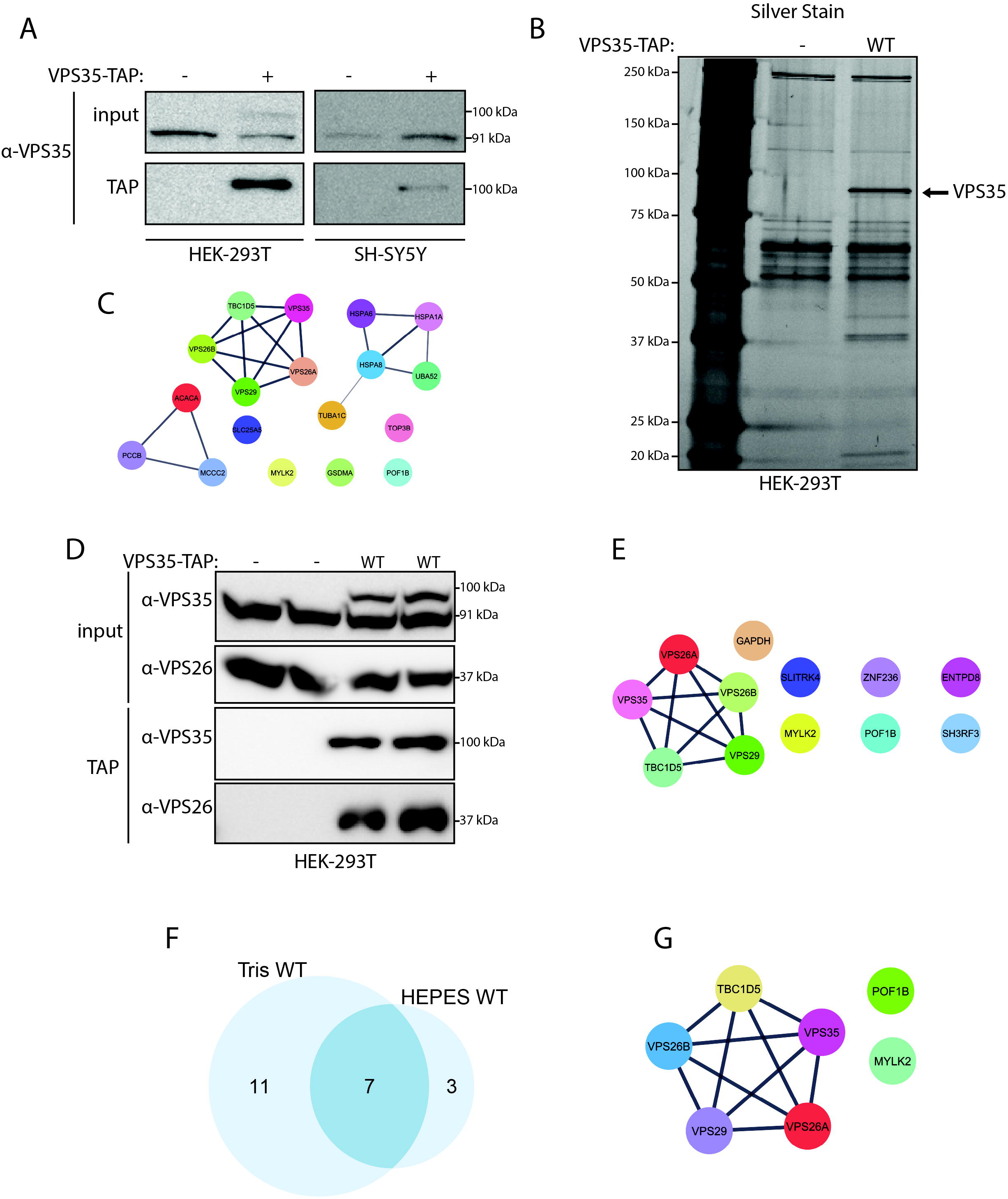
Tandem Affinity Purification (TAP) of VPS35 in HEK-293T cells using Tris– or HEPES-based buffers. (**A**) HEK-293T or SH-SY5Y cells expressing TAP-tagged WT VPS35 were subjected to TAP methodology and Western blot analysis. Inputs and TAP fractions were probed with anti-VPS35 antibody. TAP purifies VPS35 more efficiently in HEK-293T cells compared to SH-SY5Y cells. (**B**) HEK-293T cells expressing TAP-tagged WT VPS35 or empty vector (EV) were subjected to TAP methodology followed by SDS-PAGE and silver staining. Endogenous VPS35 is observed slightly below VPS35-TAP band at ∼100 kDa. (**C**) STRING diagram of interacting proteins identified by LC-MS/MS analysis of WT VPS35-TAP. (**D**) HEK-293T cells expressing TAP-tagged WT VPS35 or EV were subjected to TAP methodology with HEPES-based buffers followed by Western blot analysis. Input and VPS35-TAP fractions were probed with anti-VPS35 or anti-VPS26 antibodies to confirm recovery of the retromer and then subjected to LC-MS/MS. (**E**) STRING diagram of interacting proteins identified by LC-MS/MS analysis of WT VPS35-TAP. Outside of the core retromer subunits, no known interacting proteins of VPS35 were identified. (**F**) Proportional Venn diagram demonstrating proteins identified in WT VPS35 TAP experiments using Tris– vs. HEPES-based buffers. (**G**) STRING diagram demonstrating the 7 proteins commonly identified between the two TAP experiments.

To test the ability of TAP to identify interactors of VPS35, we subjected empty vector and TAP-tagged VPS35 pulldown fractions derived from HEK-293T cells to SDS-PAGE followed by silver staining and mass spectrometry analysis. Compared to an empty vector control, silver staining reveals TAP methodology successfully pulls down VPS35 (not present in the empty vector control) and other unknown proteins (**Figure 1B**). Mass spectrometry analysis further confirms this by identifying known core components of the retromer, including VPS26A, VPS26B, VPS29, and TBC1D5, as well as 13 other proteins not previously connected to the retromer (**Figure 1C**). To be considered a positive interactor in this experiment and all subsequent experiments, peptide intensity values for any given protein had to either be absent in the control condition or at least 4-fold enriched relative to the control.

While the identification of core components of the retromer served to validate that TAP can effectively pull-down known interacting proteins of VPS35, the low number of total recovered proteins suggested that further optimization was needed. First, we changed all TAP buffers from Tris-based buffers to HEPES-based buffers, as many papers exploring VPS35 biology in mammalian cells use HEPES-based buffers (14–16). Using HEPES-based buffers, we were able to purify TAP-tagged VPS35 and the core components of the retromer, however we were unable to improve the number of copurified proteins (**Figure 1D, E**). Interestingly, apart from the core components of the retromer, each method (Tris– vs. HEPES-based) identified novel and distinct interactions with VPS35, suggesting that the stability of retromer-interacting proteins may be affected by buffer components. When comparing Tris– vs. HEPES-based buffers, 7 proteins were commonly identified by both experimental paradigms (**Figure 1F**). Of those 7 proteins, four of them are core components of the retromer (VPS35, VPS26A, VPS26B, and VPS29) and one is a known interacting protein of the retromer (TBC1D5, a GTPase-activating protein for Rab7) (**Figure 1G**). The remaining two novel interacting proteins are involved in the cytoskeleton, MYLK2, a myosin light chain kinase, and POF1B, a protein that binds to non-muscle actin filaments (**Figure 1G**).

### Reversible protein crosslinking TAP to improve isolation of VPS35 protein interactors

Although TAP has several advantages mentioned above, we found that the two subsequent washing steps were likely too stringent to preserve labile protein interactions. With that in mind, we next tested whether a reversible chemical crosslinker, dithiobis(succinimidyl propionate) (DSP), could aid in increasing the recovery of protein interactors of VPS35 in our TAP assays. DSP is a cell-permeable crosslinker with a 12 Å spacer-arm that reacts with primary amines to crosslink lysine residues of interacting proteins and can be reversed by reducing agents (17).

To test DSP-TAP for crosslinking efficiency, reversibility, and the ability to preserve VPS35 binding to both TAP affinity resins, HEK-293T cells expressing TAP-tagged WT VPS35 or empty vector were treated with increasing concentrations of DSP followed by cell lysis and subsequent TAP. DSP-TAP fractions and input samples (reducing and non-reducing) were analyzed by Western blotting (**Figure 2A**). Using antibodies against VPS35 and β-actin, we first evaluated the non-reducing (2x Sample Buffer without DTT or β-Mercaptoethanol) and reducing (2x Sample Buffer with DTT) input samples for dose-dependent reduction (non-reducing) and recovery (reducing) of VPS35 or actin upon DSP treatment followed by treatment with reducing agents. Indeed, actin demonstrates a dose-dependent reduction under non-reducing conditions that was restored with reducing conditions. This suggests that DSP both efficiently crosslinks actin, causing high molecular weight species that do not readily enter SDS-PAGE gels, and that reducing agents successfully reverse this crosslinking (**Figure 2A**). In contrast, endogenous VPS35 levels are not diminished by DSP treatment under non-reducing or reducing conditions. Additionally, when treated with DSP, TAP-tagged VPS35 was able to bind the streptavidin resin at all concentrations tested, however 0.3 mM DSP was the only concentration in which TAP-tagged VPS35 was recovered using calmodulin affinity resin (**Figure 2A**), implying that DSP crosslinking might alter the calmodulin-binding peptide sequence within the TAP tag. For this reason, all subsequent DSP-TAP assays were performed using 0.3 mM DSP.

**Figure 2.**
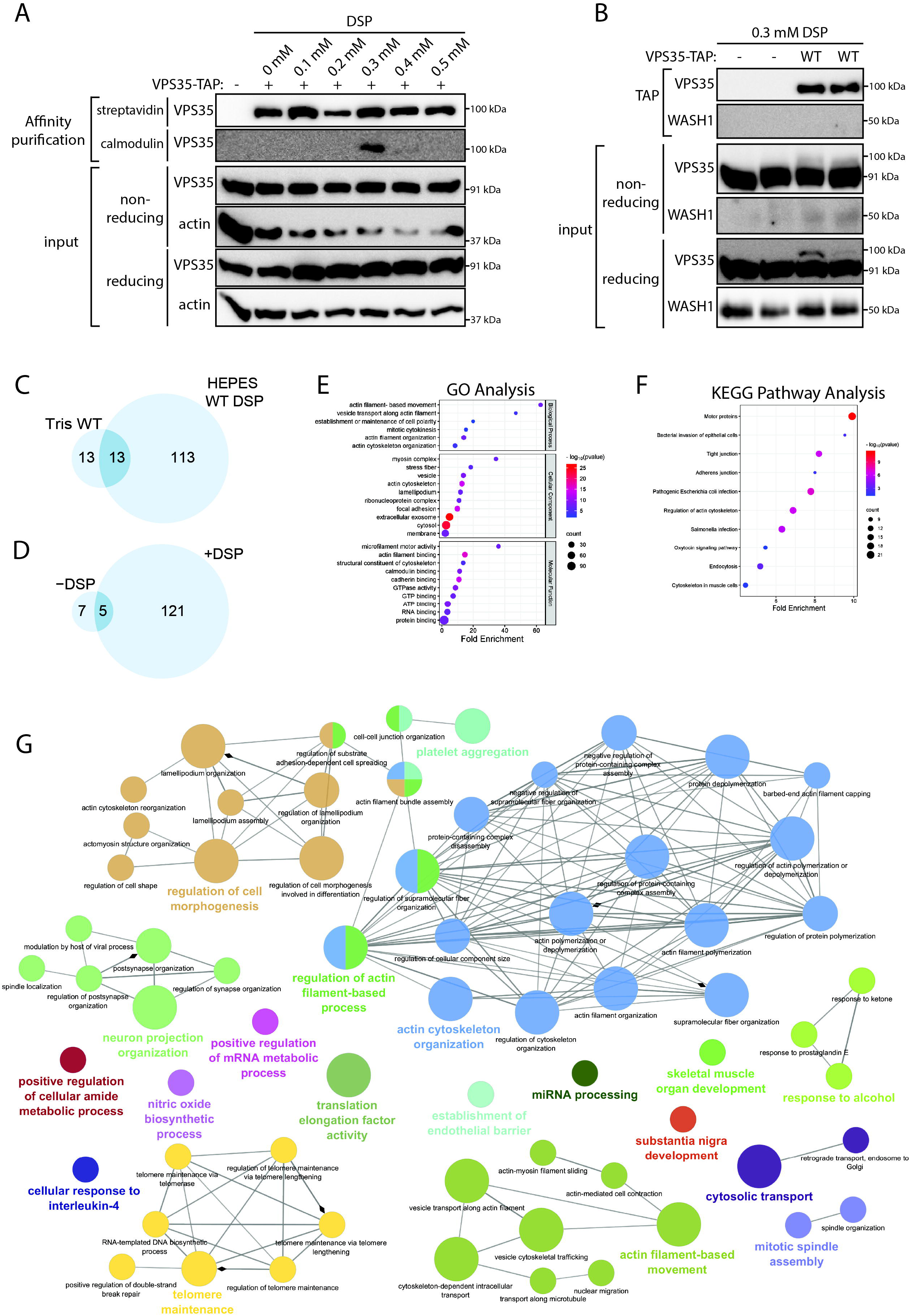
TAP of VPS35 in HEK-293T cells treated with a reversible protein crosslinker. (**A**) HEK-293T cells expressing TAP-tagged WT VPS35 treated with increasing concentrations of DSP were subjected to affinity purification with either streptavidin or calmodulin resin followed by Western blot analysis under reducing or non-reducing conditions. Input and purified fractions were probed with anti-VPS35 or anti-actin antibodies. Only 0.3 mM DSP treatment preserved binding of TAP-tagged VPS35 to both streptavidin and calmodulin resins. (B) HEK-293T cells expressing TAP-tagged VPS35 treated with 0.3 mM DSP were subjected to TAP methodology followed by Western blot analysis under reducing or non-reducing conditions. Input and TAP fractions were probed with anti-VPS35 or anti-WASH1 antibodies. WT VPS35-TAP eluate was subjected to LC-MS/MS analysis. (**C**) Proportional Venn diagram showing the common interacting proteins of WT VPS35 identified in Tris-based alone vs HEPES-based with DSP TAP experiments. (**D**) Proportional Venn diagram demonstrating proteins identified in HEPES-based WT VPS35-TAP assays with or without DSP treatment. The addition of reversible cross-linking greatly increased the number of interacting proteins. (**E**, **F**) GO and KEGG pathway analysis using the functional annotation tool DAVID of VPS35-interacting proteins identified using DSP-treated TAP assays in HEK-293T cells overexpressing TAP-tagged VPS35-WT. Top 10 GO terms for each category and top 10 KEGG Pathway terms, based on Bonferroni-corrected p-value and fold enrichment, are displayed. Bubble plots generated using SRplot. (**G**) Functional enrichment analysis using the Cytoscape plug-in ClueGo with GO biological process terms of proteins found in DSP-treated WT VPS35 TAP assays. Pathways with a p-value of <0.05 and a Kappa score of 0.4 were mapped. Two-sided hypergeometric statistical analysis test with Bonferroni step-down p-value correction was used. Node size corresponds to p-value.

To test if treatment with DSP increases the yield of VPS35 interacting proteins in our TAP assays, HEK-293T cells expressing TAP-tagged WT VPS35 or empty vector were treated with 0.3 mM DSP before being subjected to TAP and validated via Western blotting. After confirming efficient TAP-VPS35 purification via Western blotting (**Figure 2B**), VPS35 TAP or empty vector samples were subjected to LC-MS/MS analysis. Comparing TAP-tagged VPS35 levels on Western blots between reducing (2x Sample Buffer with DTT) and non-reducing (2x Sample Buffer without DTT or β-Mercaptoethanol) input conditions, we demonstrate the absence of TAP-tagged VPS35 in the non-reducing condition, with recovery in the reducing condition (**Figure 2B**). We show that using DSP-TAP in HEK-293T cells successfully purifies WT VPS35, in addition to 125 other interacting proteins, including the retromer components VPS26A, VPS26B, and VPS29 (**Table 1**, **Figure 2C**). We also show that using DSP as a reversible crosslinker successfully increased the number of copurified proteins in the TAP assays, especially compared to experiments done with Tris-based buffers or without DSP crosslinking (**Figure 2C, 2D**).

Interestingly, we were unable to detect an interaction between VPS35 and any components of the WASH complex, which has been suggested to have impaired binding to D620N VPS35 (**Figure 2B**, **Table 1**) (14). Using the Cytoscape plug-in ClueGo (18), we performed functional enrichment analysis of the WT VPS35 protein interactome from DSP-treated TAP assays and find an enrichment of proteins associated with cytosolic transport, actin cytoskeleton organization, and neuron projection organization, all of which have been previously shown to directly involve VPS35 and/or the retromer (**Figure 2G**) (7,8,10,11,19). We also find an enrichment in proteins associated with cellular processes that have not previously been associated with VPS35 and the retromer, such as telomere maintenance, representing potential novel pathways involving the retromer (**Figure 2G**). Additionally, using the functional annotation tool DAVID (20,21), we performed gene ontology (GO) and KEGG pathway analyses of the 125 proteins identified to interact with WT VPS35 (**Figure 2E, 2F**). GO biological pathway analysis highly correlated with ClueGo functional enrichment results and reveals significant enrichment of proteins involved in actin-related pathways and vesicular transport (**Figure 2E**). As expected, GO cellular compartment analysis shows an enrichment for proteins located at the membrane and cytosol, with additional enrichment in the extracellular exosome (**Figure 2E**). GO molecular function analysis shows an expected enrichment of proteins involved in protein binding and actin cytoskeleton activity (**Figure 2E**). Given the known function of VPS35 and the retromer in retrograde cellular transport, KEGG pathway analysis shows an enrichment of motor proteins and actin cytoskeleton regulators (**Figure 2F**).

### Traditional co-IP with reversible protein crosslinking

Given the low number of interactors identified in our TAP assays, even with extensive experimental optimization, we suspect that TAP, although detergent-free, may be unable to isolate labile protein interactions with the retromer, given its two consecutive purification steps. Importantly, our TAP assays were unable to recover any components of the WASH complex, which is a well-characterized accessory complex to the retromer. This could be due to stringent purification steps or to the low levels of TAP-tagged VPS35 at steady-state. Additionally, we cannot rule out interference of the TAP tag itself with WASH complex binding at the C-terminus of VPS35. Due to these caveats, we decided that TAP methodology was likely not optimal for the identification of differential protein interactors with the D620N mutation. However, it is important to not understate the stable interactions that our DSP-TAP assays identified as these may represent novel pathways in which VPS35 and the retromer are involved. Further investigation into these pathways is important in understanding retromer biology and could potentially lead the field in new directions.

We next wanted to determine if traditional co-immunoprecipitation (co-IP) assays combined with chemical crosslinker treatment could improve the number of identified protein interactors of VPS35 and recover components of the WASH complex. To test the efficiency of a co-IP assay with cross-linker treatment, HEK-293T cells expressing V5-tagged WT or D620N VPS35, or empty vector, were subjected to DSP treatment and subsequent IP with anti-V5 antibody. A small portion of the input and IP were subjected to Western blot analysis to confirm the equivalent pulldown of V5-tagged VPS35 variants (**Figure 3A**). LC-MS/MS analysis identifies 303 interacting proteins between WT and D620N VPS35 conditions, 297 of which are shared between the two conditions, highlighting high similarity between the interactomes of both WT and D620N VPS35 (**Figure 3E**, **Table 2**). Peptide intensities were normalized to the average intensity of VPS35 among all samples, excluding the empty vector control. To be considered an authentic interactor of VPS35, peptide intensity values for any given protein had to be absent in the control condition or at least 4-fold enriched relative to the control. If a given protein was present in all conditions, but only 4-fold enriched in one, it was considered absent in the other. For example, PGAM1 is enriched an average of 5.9-fold above control in the WT condition, but only 2.8-fold enriched above control in the D620N condition. In our Venn diagram, PGAM1 represents one of the 6 proteins found only in the WT condition (**Figure 3E**).

**Figure 3.**
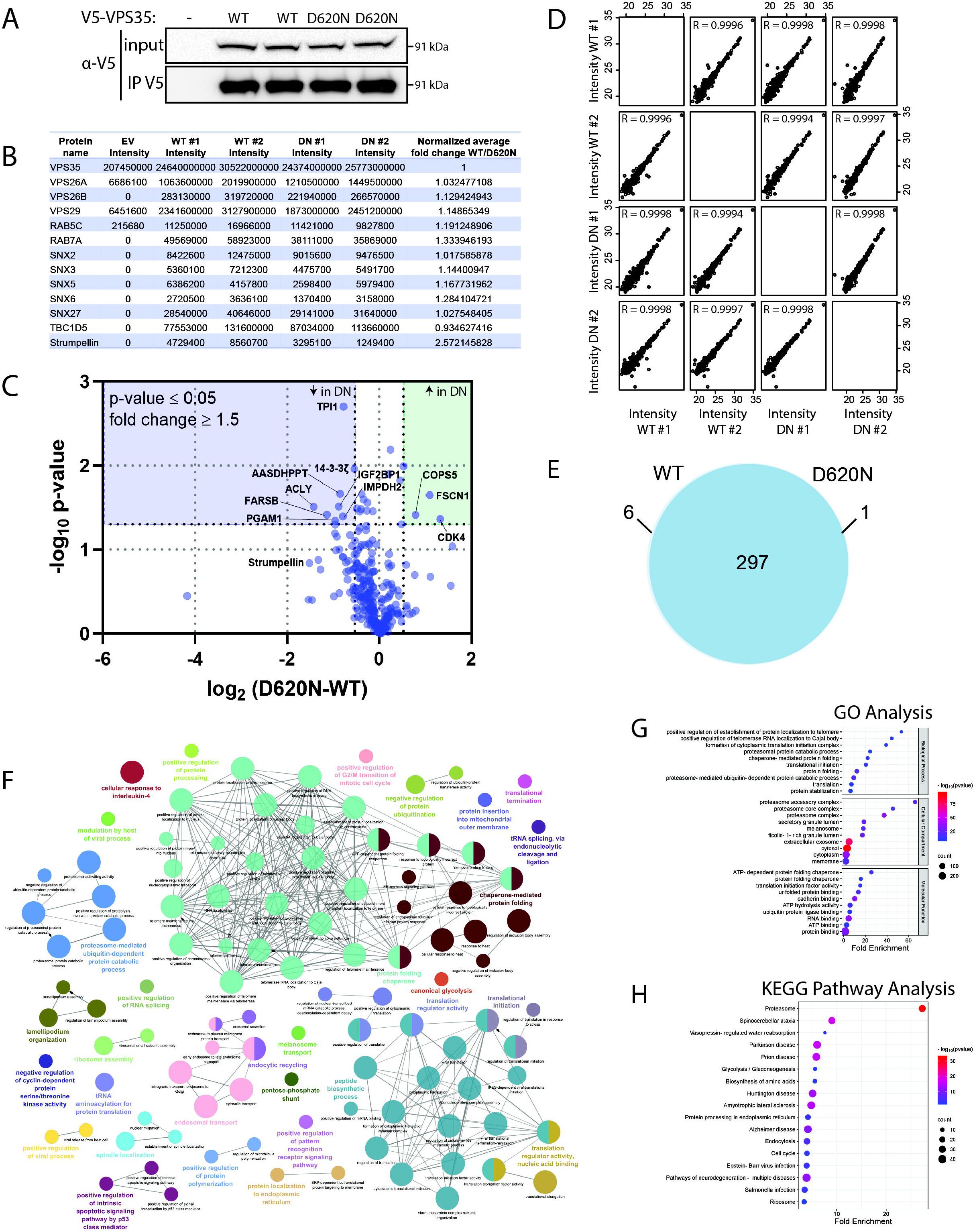
Co-IP of VPS35 in HEK-293T cells treated with a reversible protein crosslinker. (**A**) HEK-293T cells expressing V5-tagged VPS35 (WT or D620N) and treated with DSP were subjected to IP with anti-V5 antibody followed by Western blot analysis. Input and IP fractions were probed with anti-V5 antibody followed by LC-MS/MS analysis. (**B**) Table highlighting proteins known to interact with VPS35 that were identified by LC-MS/MS, indicating peptide intensity values for each interactor across conditions and replicates. (**C**) Volcano plot of LC-MS/MS data plotted as the log_2_ fold change of WT vs D620N VPS35 co-IP samples against the –log_10_ p-value. Significance was considered for p-values ≤ 0.05 and fold changes greater than 1.5 (*n* = 2/group). (**D**) Correlation analysis comparing the log_2_ transformed intensity values of proteins found in each condition/replicate compared with all other conditions. All samples were highly correlated with each other (R > 0.99). (**E**) Proportional Venn diagram demonstrating proteins identified in V5-tagged WT vs D620N VPS35 co-IP experiments. (**F**) Functional enrichment analysis using the Cytoscape plug-in ClueGo with GO biological process terms of proteins found in DSP-treated V5-tagged WT VPS35 co-IP assays. Pathways with a p-value of <0.05 and a Kappa score of 0.4 were mapped. Two-sided hypergeometric statistical analysis test with Bonferroni step-down p-value correction was used. Node size corresponds to p-value. (**G**, **H**) GO and KEGG pathway analysis using the functional annotation tool DAVID of VPS35 interacting proteins identified by LC-MS/MS from HEK-293T cells overexpressing V5-tagged WT VPS35 treated with DSP. Top 10 GO terms for each category and top 20 KEGG Pathway terms, based on Bonferroni-corrected p-value and fold enrichment, were displayed. Bubble plots generated using SRplot.

In addition to the core components of the retromer complex (VPS35, VPS26A, VPS26B, and VPS29), we are able to recover several known accessory components of the retromer including TBC1D5, Strumpellin, sorting nexins (SNX2, 3, 5, 6 and 27), and Rab proteins (Rab5c, Rab7a) (**Figure 3B**). Strumpellin, a component of the pentameric WASH complex, is reduced on average by 2.5-fold in D620N VPS35 compared to WT VPS35 samples. This is the only known interactor of the retromer that shows an average fold-change greater than 2 in this experiment (**Figure 3B**). When normalized peptide intensities are plotted as a volcano plot, where the log_2_ fold change is plotted against the –log_10_ p-value, Strumpellin does not reach our significance threshold (p-value ≤ 0.05 and fold change ≥ 1.5), however, we partially replicated previous findings by McGough *et. al*. (22) suggesting a decrease in WASH complex binding to D620N VPS35 (**Figure 3B, 3C**). Consistent with previous data from our group and others suggesting retromer function remains largely intact with the D620N mutation, correlation analysis comparing the log_2_-transformed intensity values between all conditions indicates a high level a similarity with R-values all above 0.99 between samples (**Figure 3D**) (14,22–24). These data suggest that the effects of the D620N mutation on VPS35 protein interactions is likely subtle.

In these co-IP assays, we identify 8 proteins with significantly decreased binding to D620N VPS35 compared to WT protein (**Figure 3C**). Three of these proteins (PGAM1, TPI1, and ACLY) are directly involved in ATP production through aerobic respiration. Specifically, PGAM1 and TPI1 are involved in glycolysis. PGAM1 is responsible for converting 3-phosphoglycerate to 2-phosphoglycerate, while TPI1 is responsible for the reversible conversion of dihydroxyacetone phosphate to glyceraldehyde-3-phosphate (25). ACLY, an enzyme present in the cytosol, catalyzes the conversion of citrate, a product of the mitochondrial citric acid cycle, to acetyl-CoA and oxaloacetate, essentially bridging carbohydrate metabolism to other biosynthetic pathways. Specifically, acetyl-CoA is an essential building block for fatty acid synthesis, cholesterol synthesis, and histone acetylation (26). Additionally, AASDHPPT and 14-3-3ζ are also suggested to be involved in parallel biosynthetic processes. L-aminoadipate-semialdehyde dehydrogenase-phosphopantetheinyl transferase (AASDHPPT) has recently been shown to be a key enzyme in mitochondrial fatty acid synthesis (mtFAS) (27). 14-3-3ζ is suggested to be indirectly involved in glycolysis through its role in canonical signaling pathways that regulate the glycolytic pathway, including PI3K/Akt and β-catenin/c-Myc (28,29). These findings suggest that VPS35 and the retromer may have a role in metabolic processes that have not been previously described. Furthermore, dysfunction of these pathways could potentially contribute to the pathogenic effects of D620N VPS35, however, further validation of these metabolic effects are now warranted.

We also identify 3 proteins that show significantly increased binding to D620N VPS35. These include COPS5, FSCN1, and CDK4, which are involved in cullin-RING ubiquitin ligase maintenance, actin cytoskeleton regulation, and cell cycle control, respectively (**Figure 3C**) (30–32). These proteins have not previously been described as interactors of VPS35 and the retromer. Furthermore, these proteins do not seem to be involved in similar pathways, unlike several of the proteins we identified to be decreased in the D620N condition. It remains unclear whether these proteins contribute to the pathogenic effects of D620N VPS35.

Similar to our DSP-TAP assays, we used the Cytoscape plug-in ClueGo (18) to perform functional enrichment analysis of the WT VPS35 protein interactome from HEK-293T cells overexpressing VPS35 and treated with the reversible protein cross-linker DSP. Like DSP-TAP, functional enrichment analysis identified several pathways in which VPS35 and the retromer are known to be involved, including endosomal transport and endocytic recycling (**Figure 3F**). Interestingly, proteins related to the actin cytoskeleton are not identified, highlighting that these interactors may only be purified under native conditions (**Figure 3F**). Functional enrichment analysis also identifies several proteins involved in chaperone-mediated protein folding, translational regulation, and peptide biosynthesis, all of which have not been directly linked to the retromer (**Figure 3F**). Consistent with functional enrichment analysis, GO biological function analysis reveals enrichment of proteins involved in protein stabilization, translation, and proteasome-mediated protein catabolism (**Figure 3G**). As expected, GO cellular compartment analysis in this model shows an enrichment for proteins located in the membrane, cytosol, and extracellular exosome (**Figure 3G**). GO molecular function analysis shows an expected enrichment in proteins involved in protein binding, but interestingly also indicates an enrichment for proteins involved in ATP and RNA binding (**Figure 3G**). Unexpectedly, KEGG pathway analysis reveals high enrichment for the proteasome pathway (**Figure 3H**), suggesting an uncharacterized role of the retromer in the recognition, sorting or removal of misfolded and damaged proteins, which is particularly interesting in the context of neurodegenerative disease.

### Viral-mediated expression of human VPS35 variants in the rat nigrostriatal pathway

To determine differential protein interactions of VPS35 variants *in vivo*, adult female Sprague Dawley rats were bilaterally injected into the substantia nigra with recombinant AAV2/6 vectors expressing V5-tagged human VPS35 (WT or D620N) and empty control virus (MCS) in opposite hemispheres. At 4 weeks post-injection, soluble hemi-brain extracts were subjected to IP with an anti-V5 antibody, followed by Western blotting and LC-MS/MS analysis (**Figure 4A**). Proteins identified in this manner represent interactors of VPS35 within the nigrostriatal pathway due to intranigral delivery of AAV-VPS35 vectors. Western blot analysis confirms the enrichment of V5-tagged VPS35 variants in the injected nigra and the co-IP of endogenous VPS26 (**Figure 4A**), confirming that human VPS35 integrates into the endogenous retromer complex. Peptide intensities detected by LC-MS/MS were normalized to the average intensity of VPS35 among all samples, excluding the MCS control. To be considered a positive interactor of VPS35, peptide intensity values for any given protein had to be absent in the control condition or at least 4-fold enriched relative to the control. As with data generated from cell lines, WT and D620N VPS35 protein interactomes share a high degree of similarity, with 290 proteins identified in both conditions (**Figure 4C**, **Table 3**). As expected, all conditions were highly correlated with each other, with the lowest R-value being 0.9967, further suggesting any effects of the D620N mutation on VPS35 function is likely to be subtle (**Figure 4B**). Importantly, we identify known retromer components for WT and D620N VPS35, including VPS26A, VPS26B, VPS29 and TBC1D5 (**Table 3**), confirming that human VPS35 successfully integrates into the endogenous retromer complex. Intriguingly, this experiment identifies a small number of significantly enriched proteins in the WT or D620N VPS35 conditions, representing putative differential interacting proteins that may play a role in retromer dysfunction underlying VPS35-induced neurodegeneration (**Figure 4D**). The proteins identified in this experiment represent the first brain-specific protein interactome of VPS35 and the retromer (**Figure 4D**). In addition to the 18 proteins with significantly decreased binding to D620N VPS35, 49 proteins completely failed to bind D620N VPS35 relative to WT protein, representing putative binding partners that may play a role in D620N VPS35-induced pathogenicity (**Figure 4D**). The most abundantly represented pathways determined by STRING analysis, using MCL clustering, among the 67 proteins either absent or depleted in the D620N VPS35 condition include the RHOBTB2 GTPase cycle, the β-catenin destruction complex, mRNA processing, and actin cytoskeleton, with only the latter representing a pathway previously linked to the function of the retromer (**Figure S1A**) (19). We also found 19 proteins with either absent or depleted binding to WT VPS35 (i.e. enriched or unique to D620N VPS35), however these 19 proteins do not significantly cluster using STRING analysis (**Figure S1B**).

**Figure 4.**
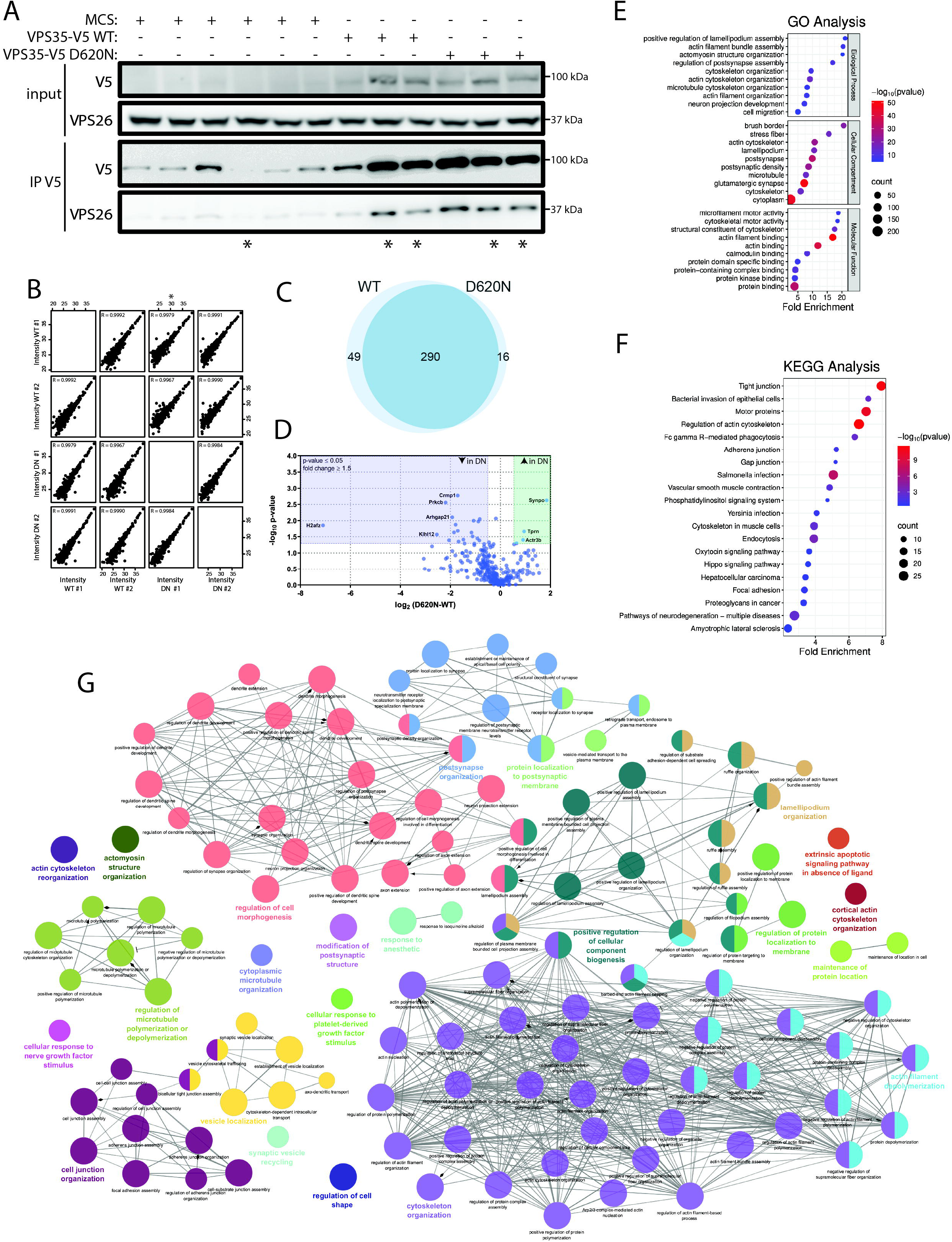
Co-IP of human VPS35 overexpressed in the rat nigrostriatal pathway. (**A**) Adult female Sprague Dawley rats with bilateral intranigral injections of AAV2/6 vectors expressing V5-tagged VPS35 (WT or D620N) in one hemisphere and control virus (MCS) in the opposite. Triton-soluble brain extracts derived from rat hemi-brains at 4 weeks were subjected to IP with anti-V5 antibody, followed by Western blot analysis with anti-V5 and anti-VPS26 antibodies. Asterisks indicate the five samples subjected to LC-MS/MS analysis (**B**) Correlation analysis comparing the log_2_ transformed intensity values of proteins found in each condition/replicate compared with all other conditions. All samples were highly correlated with each other (R > 0.99). (**C**) Proportional Venn diagram demonstrating interacting proteins identified in the nigrostriatal pathway of rats injected with V5-tagged WT vs D620N VPS35. (**D**) Volcano plot of LC-MS/MS data plotted as the log_2_ fold change of WT vs D620N VPS35 injected animals against the –log_10_ p-value. Significance was considered for p-values ≤ 0.05 and fold changes greater than 1.5 (*n* = 2/group). (**E**, **F**) GO and KEGG pathway analysis using the functional annotation tool DAVID of VPS35 interacting proteins identified by LC-MS/MS. Top 10 GO terms for each category and top 20 KEGG Pathway terms, based on Bonferroni-corrected p-value and fold enrichment, were displayed. Bubble plots generated using SRplot. (**G**) Functional enrichment analysis using the Cytoscape plug-in ClueGo with GO biological process terms of proteins found in the nigrostriatal pathway of AAV-V5-VPS35 injected rats. Pathways with a p-value of <0.05 and a Kappa score of 0.4 were mapped. Two-sided hypergeometric statistical analysis test with Bonferroni step-down p-value correction was used. Node size corresponds to p-value.

Importantly, we demonstrate the first brain-specific protein interactome for VPS35 and the retromer, highlighting putative pathways that may be important specifically in the nigrostriatal pathway. With that in mind, we performed functional enrichment analysis on proteins identified in the WT VPS35 condition to determine retromer-mediated pathways specific to the brain (**Figure 4E-G**). As expected, proteins involved in actin cytoskeleton and microtubule organization, postsynaptic membrane organization, and vesicle localization are enriched and have previously been linked to retromer function (8,19). Unexpectedly, functional enrichment analysis identifies putative pathways that may require the retromer but have not been previously described. These include regulation of cell morphogenesis, cellular component biogenesis, and cell junction organization (**Figure 4E**).

### Brain-specific VPS35 interactome analysis in *D620N VPS35* knockin mice

To date, protein interactome studies of WT and D620N VPS35 have relied on either protein overexpression in cell culture models or conditional knockout mouse models of *VPS35*, which do not sufficiently recapitulate neurodegeneration (22,33–36) Accordingly, we performed interactome and global proteomic analysis in *D620N VPS35* knockin (KI) mice (37,38). Importantly, these mice express WT or D620N VPS35 at endogenous levels, making this model physiologically relevant to disease, compared to overexpression or knockout models. We first performed co-IP assays with Triton-soluble hemi-brain extracts, where one half of the brain was used for an anti-VPS35 antibody IP and the remaining half for an isotype-matched antibody IP as a control. For this first assay, WT, heterozygous D620N KI, and homozygous D620N KI mice were subjected to IP, however, only homozygous animals were analyzed via LC-MS/MS. Western blot analysis demonstrates the specific immuno-purification of VPS35 from soluble hemi-brain extracts and its co-IP with the core-selective complex subunit VPS26 (**Figure 5A**).

**Figure 5.**
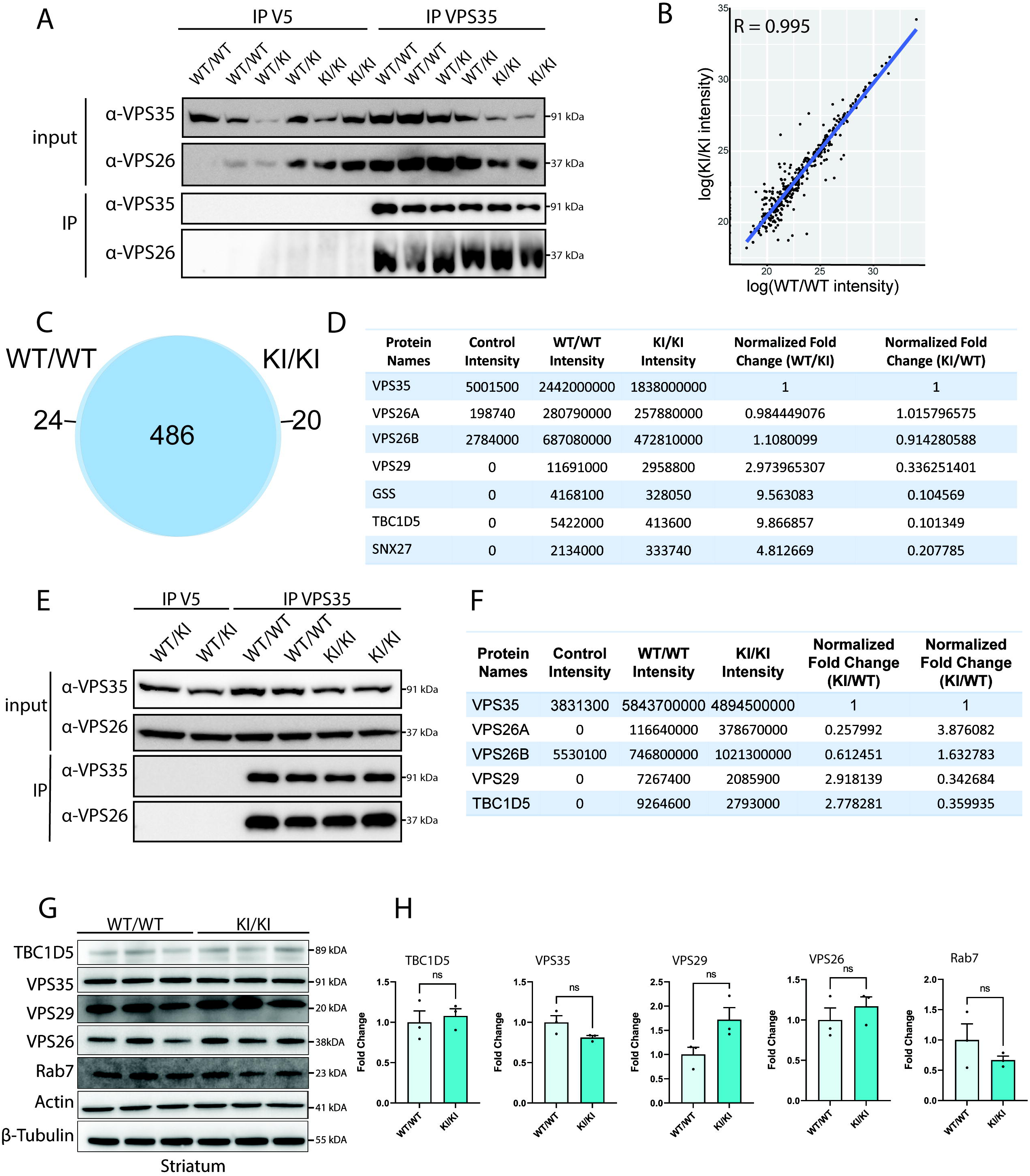
Co-IP of VPS35 in hemi-brain or striatum from *D620N VPS35* knockin mice. (**A**) Soluble hemi-brain extracts from 3-4 month-old WT, heterozygous and homozygous KI mice were subjected to IP with anti-VPS35 antibody (or isotype-matched control anti-V5 IgG) followed by Western blot analysis. Input and IP fractions were probed with anti-VPS35 and anti-VPS26 antibodies, prior to subjecting VPS35 IP samples to LC-MS/MS analysis. (**B**) Correlation analysis of VPS35-interacting proteins from WT vs homozygous *D620N VPS35* KI reveal a high degree of correlation (R = 0.995). (**C**) Proportional Venn diagram demonstrating the strong overlap between interacting proteins identified in WT and homozygous *D620N VPS35* KI brains. (**D**) Table highlighting peptide intensities across conditions for core retromer subunits, and three interacting proteins depleted in *D620N VPS35* KI brain. (**E**) Soluble striatal extracts from 3-4 month-old WT and KI mice were subjected to IP with anti-VPS35 antibody (or isotype-matched control anti-V5 IgG) followed by Western blot analysis. Input and IP fractions were probed with anti-VPS35 and anti-VPS26 antibodies, prior to subjecting VPS35 IP samples to LC-MS/MS analysis. (**F**) Table outlining peptide intensities across conditions for core retromer subunits in striatal tissue of *D620N VPS35* KI mice. (**G**) Triton-soluble fractions from striatum of adult WT or *D620N VPS35* KI mice were subjected to Western blot analysis to monitor steady-state levels of core retromer subunits (VPS35, VPS26, VPS29), Rab7 and TBC1D5. (**H**) Graphs indicate densitometric analysis of protein levels normalized to actin or β-tubulin and expressed as fold-change compared to WT mice (mean ± SEM, *n* = 3 mice/group). Data are not significant (*P*>0.05) by unpaired, two-tailed Student’s *t*-test.

We next subjected one WT and one homozygous *D620N VPS35* KI IP to LC-MS/MS analysis. The VPS35 interactome of WT and D620N VPS35 KI hemi-brains share a high degree of similarity, as shown by correlation analysis and an R-value of 0.995 (**Figure 5B**). This observation is consistent with the data generated using overexpression models and further suggests that the D620N mutation has a subtle effect on the VPS35 interactome. After normalizing peptide intensities to the average level of VPS35 between conditions, 486 interacting proteins are identified across both genotypes, with 24 proteins found to only bind WT VPS35, and 20 found to only bind D620N VPS35 (**Figure 5C**, **Table 4**). Our analysis reveals the successful co-IP of cargo-selective complex components, including VPS35, VPS26A, VPS26B, and VPS29 (**Figure 5D**). Interestingly, VPS26B is detected at 2.1-fold increased levels over VPS26A, suggesting that VPS26B may have a brain-specific cargo sorting role and/or is more abundant in brain tissue. Notably, several proteins are found to be enriched or depleted in binding to D620N VPS35. Specifically, VPS29, GSS, TBC1D5, and SNX27 show reduced binding to D620N VPS35, three of which (VPS29, TBC1D5, and SNX27) are known to directly interact with the retromer (**Figure 5D**). Interestingly, the D620N VPS35 mutation does not impact these interactions in overexpression models in human cells or rat brain (**Figures 1-4**). This could potentially be explained by VPS35 overexpression altering the stoichiometry of the endogenous retromer that masks the effects of the D620N mutation, and/or these disruptions could be specific to brain tissue that would preclude them from being identified in HEK-293T cells. We are unable to detect an interaction of VPS35 with other sorting nexins (SNX1, 2, 3, 5, 6), WASH complex components, or known retromer cargo in rodent brain tissue. We have previously shown that WASH1 and Fam21 are abundant in whole brain extracts (38), suggesting that the WASH complex is a labile interactor and/or may not be required for retromer-mediated cargo sorting in the brain.

To further explore the brain-specific VPS35 protein interactome in a PD-relevant brain region, we performed similar co-IP assays on striatal extracts from WT and homozygous *D620N VPS35* KI mice with an anti-VPS35 antibody. Striatum from heterozygous KI animals was used for an isotype-matched antibody control IP. Western blot analysis of co-IP samples reveals equivalent pulldown of VPS35 between genotypes and confirms the known interaction with VPS26 (**Figure 5E**). As with the VPS35 co-IP from hemi-brain, we observe a ∼3-fold reduction in the binding of D620N VPS35 to VPS29 and TBC1D5, confirming our findings in a region of the brain affected in PD (**Figure 5F**, **Table 5**). To determine if the decrease in VPS29 and/or TBC1D5 binding to D620N VPS35 was simply due to a decrease in protein steady-state levels, we assessed striatal extracts by Western blot and densitometry analysis (**Figure 5G**, **H**). We confirm that there are no significant changes in the levels of core-selective complex subunits (VPS35, VPS26, VPS29), TBC1D5 or Rab7 in the striatum of *D620N* KI mice (**Figure 5H**) (16).

### Functional enrichment analysis of endogenous VPS35 protein interactors in the brain

While we have identified putative interacting proteins of VPS35 that may be disrupted by the D620N mutation and contribute to neurodegeneration, we have also identified the first brain-specific protein interactome of endogenous WT VPS35. Importantly, functional enrichment analysis of VPS35 interacting proteins from mouse whole brain extracts reveals several cellular pathways, both well-described and novel, related to VPS35 and the retromer (**Figure 6A**). Expected pathways based on the known function of the retromer include cytosolic transport, dopamine metabolism, protein localization, neuronal death, regulation of protein-containing complex assembly, and postsynaptic neurotransmitter regulation (**Figure 6A**) (8,11). Uncharacterized pathways enriched in the VPS35 protein interactome include several metabolic processes (glycolysis, peptide metabolism, DNA metabolism, glutathione metabolism), cellular detoxification, ribonucleoprotein complex organization, protein folding chaperone, and proteasomal protein catabolism (**Figure 6A**). These represent putative retromer-mediated pathways that could be specific to the brain. Interestingly, GO biological process analysis reveals the highest enrichment in proteins involved in glycolysis and the mitochondrial citric acid cycle (**Figure 6B**). Proteins from both pathways are also identified in DSP-treated co-IP assays in HEK-293T cells and specifically show reduced binding to D620N VPS35 (**Figure 3F-H**), although the same trends are not observed in hemi-brain co-IP assays from rat brain (**Figure 4E-G**). GO molecular function analysis reveals enrichment for proteins involved in many forms of protein binding, whereas GO cellular component analysis indicates an enrichment for proteins located in the cytoplasm, synapse, and nucleus (**Figure 6B**). Consistent with functional enrichment analysis, KEGG pathway analysis shows an enrichment of proteins involved in many forms of metabolism as well as endocytosis, cytoskeletal regulation, and pathways of neurodegeneration (**Figure 6C**). Importantly, this study reveals a first glimpse at the brain-specific protein interactome of VPS35, identifying putative pathways involved in retromer biology. It will be important for future work to dissect these pathways and determine the exact role of the retromer.

**Figure 6.**
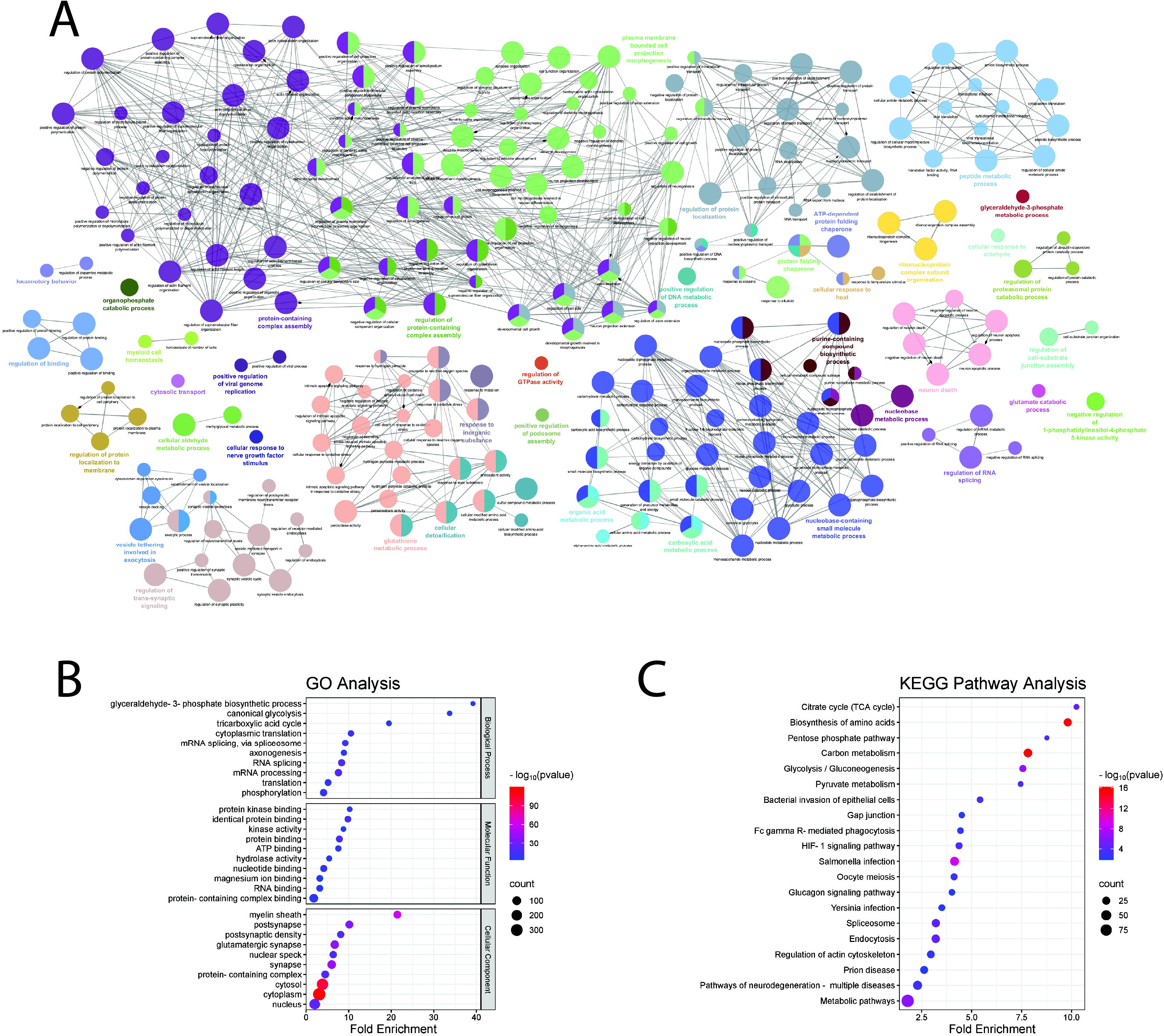
Functional enrichment analysis of VPS35-interacting proteins in brains of WT mice. (**A**) Functional enrichment analysis using the Cytoscape plug-in ClueGo with GO biological process terms of VPS35-interacting proteins identified in soluble hemi-brain extracts of WT mice. Pathways with a p-value of <0.05 and a Kappa score of 0.4 were mapped. Two-sided hypergeometric statistical analysis test with Bonferroni step-down p-value correction was used. Node size corresponds to p-value. (**B**, **C**) GO and KEGG pathway analysis using the functional annotation tool DAVID of VPS35-interacting proteins from soluble hemi-brain fractions of WT mice identified by LC-MS/MS. Top 10 GO terms for each category and top 20 KEGG Pathway terms, based on Bonferroni-corrected p-value and fold enrichment, were displayed. Bubble plots generated using SRplot.

### Comparison of brain-specific VPS35 co-IP approaches

We next wanted to determine the similarities and differences between the VPS35 protein interactome among our brain co-IP assays. When comparing our three brain-specific co-IP assays, intensity levels of VPS35 are within the same order of magnitude among all samples, with levels between 1.8 to 5.8 x 10^9^ (**Figure 7A**). VPS35 is most abundant in mouse striatum, ∼2.5-fold above the average levels of VPS35 in the rat substantia nigra and mouse whole brain assays. The elevated abundance of recovered VPS35 protein does not equate to a higher number of identified interacting proteins, as the mouse whole brain co-IP recovers more proteins than the mouse striatum or rat substantia nigra co-IPs (**Figure 7A**). When comparing proteins that meet our threshold criteria (absent or at least 4-fold enriched relative to control) in all three WT conditions, we identify 38 proteins that are common among all assays (**Figure 7B**). Using the Cytoscape plug-in ClueGo (18), we performed functional enrichment analysis on these 38 common interacting proteins. As expected, enriched pathways include cytosolic transport, protein complex and autophagosome assembly, and actin regulation (**Figure 7C**). Similarly, functional enrichment analysis of the 43 proteins commonly identified among D620N VPS35 conditions indicates similarly enriched pathways compared to WT VPS35, including cytosolic transport, protein complex assembly, and microtubule regulation (**Figure 7D-E**).

**Figure 7.**
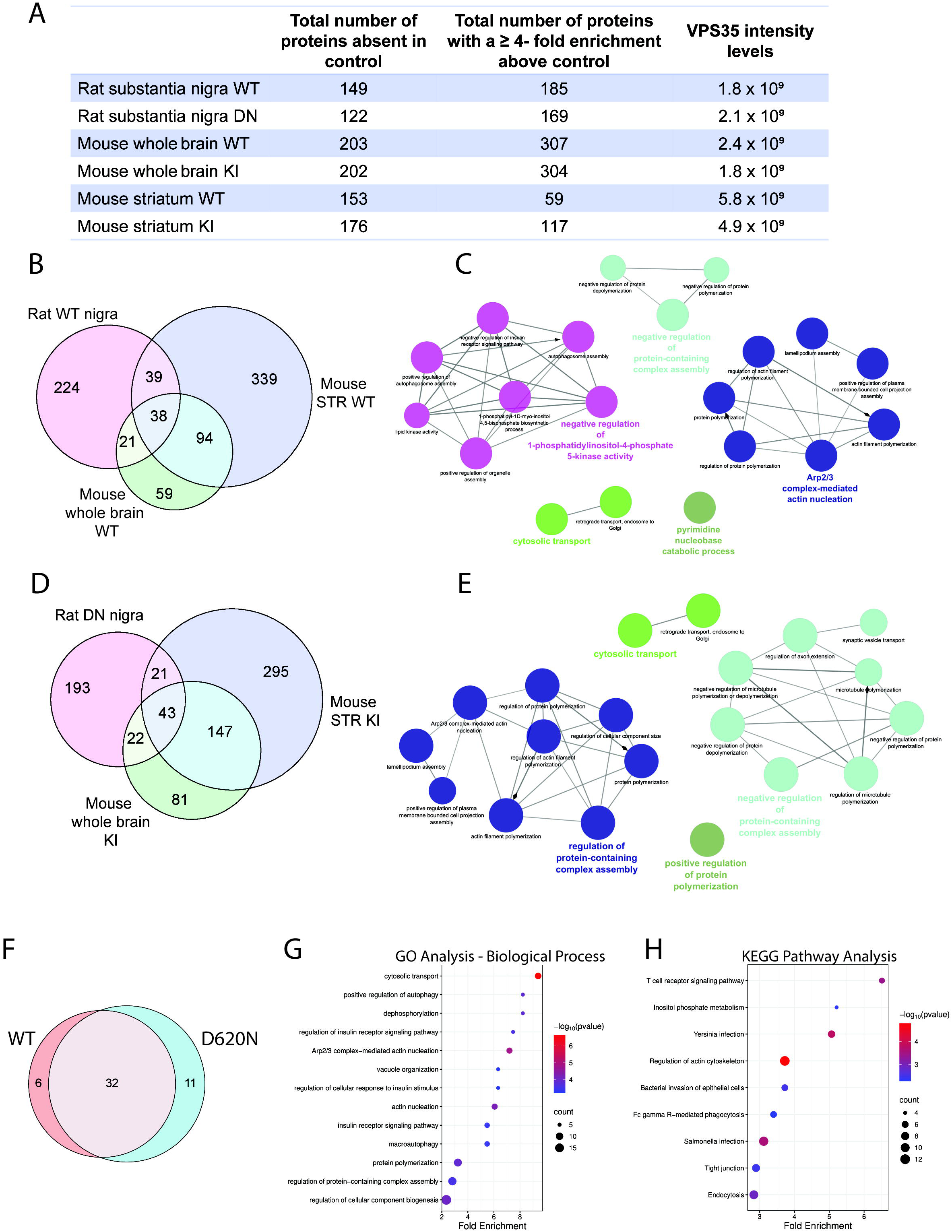
Comparison of VPS35-interacting proteins identified using different brain co-IP approaches. **(A)** Table summarizing the number of interacting proteins detected and corresponding VPS35 peptide intensity levels in three brain co-IP experiments: rat substantia nigra injected with AAV-VPS35 (WT or D620N), whole brain from WT or *D620N VPS35* KI mice, and striatum (STR) from WT or *D620N VPS35* KI mice. (**B**, **D**) Proportional Venn diagrams demonstrating the overlap between WT (**B**) or D620N VPS35 (**D**) conditions for three brain co-IP experiments. 38 proteins were commonly identified across all three WT experiments, and 43 proteins across all three D620N experiments. (**C**, **E**) Functional enrichment analysis using the Cytoscape plug-in ClueGo with GO biological process terms of 38 WT VPS35-interacting proteins (**C**) or 43 D620N VPS35-interacting proteins (**E**) commonly identified across the three brain co-IP experiments. Pathways with a p-value of <0.05 and a Kappa score of 0.4 were mapped. Two-sided hypergeometric statistical analysis test with Bonferroni step-down p-value correction was used. Node size corresponds to p-value. (**F**) Proportional Venn diagram comparing the 38 or 43 VPS35-interacting proteins commonly identified across all three WT or D620N experiments, respectively. 32 proteins were common between WT and D620N VPS35. (**G**, **H**) GO and KEGG pathway overrepresentation analysis of the model agnostic 49 VPS35-interacting proteins shown in (**F**) using the “enrichGO” and “enrichKEGG” functions in the R package “clusterProfiler.” Significantly enriched pathways based on Bonferroni-corrected p-value and fold enrichment, were displayed. Bubble plots generated using SRplot.

When comparing the 38 and 43 proteins commonly found across WT and D620N VPS35 brain co-IP assays, respectively, we find 32 proteins common to both conditions, 6 proteins found only with WT VPS35, and 11 proteins found only with D620N VPS35 (**Figure 7F**). Next, we sought to determine which, if any, pathways are overrepresented in our three brain co-IP experimental paradigms. We performed overrepresentation analysis (ORA) using a subset of proteins commonly found in WT or D620N conditions, specifically the 49 proteins represented in **Figure 7F** (39). Our pre-defined protein list included 870 interacting proteins that meet our significance criteria in at least one of the conditions of our three brain co-IP assays (**Figure 7B, D**). When comparing the subset of 49 proteins commonly found in WT or D620N conditions to our pre-defined list of 870 proteins, ORA indicates an enrichment of GO terms associated with biological process, as well as several KEGG pathway analysis terms (**Figure 7G, H**). These pathways represent known and putative functional roles of VPS35 and the retromer across two different species (rat, mouse) and three different brain regions (nigra, striatum, whole brain). GO biological process terms include pathways in which the retromer has well-defined roles, including cytosolic transport, regulation of protein-containing complex assembly, and vacuole organization (**Figure 7G**) (7,8,11,19). ORA of GO terms additionally provides insight into how VPS35 and the retromer may be critical for cellular homeostasis in brain tissue. ORA of KEGG pathway analysis demonstrates that VPS35 and the retromer have a ubiquitous role in endocytosis in the brain, as well as involvement in several immune response pathways (**Figure 7H**). Taken together, we demonstrate that VPS35 and the retromer play a critical role in the brain and suggest several pathways that may be required for brain homeostasis, regardless of specific brain region or cell type.

### Validation of TBC1D5 as a differential interactor of D620N VPS35

One consistent finding in both hemi-brain and striatal co-IP assays from WT and D620N VPS35 KI mice is the reduced binding of VPS29 and TBC1D5 to D620N VPS35 relative to WT protein. Previous studies have suggested that the interaction of the core components of the retromer are not altered by the D620N mutation, however, these studies relied on overexpression in cultured cells (22,24,33,36). We aimed to validate diminished binding to TBC1D5 in co-IP assays with anti-VPS35 antibody on Triton-soluble striatal extracts from WT and homozygous *D620N VPS35* KI mice. Striatal tissue from heterozygous KI mice were used as an isotype-matched antibody control IP (anti-V5). Western blot analysis reveals decreased pulldown of TBC1D5 in D620N KI compared to WT mice, thereby validating our LC-MS/MS data (**Figure 8A**). Next, we elected to validate TBC1D5 binding capacity to human VPS35 variants in another system. HEK-293T cells overexpressing V5-tagged VPS35 variants were subjected to IP with anti-V5 antibody, and input and IP fractions were subjected to Western blot analysis with anti-TBC1D5 and anti-V5 antibodies. We demonstrate that endogenous TBC1D5 can interact with WT and D620N VPS35, however, the D620N mutation significantly decreases binding to TBC1D5 (**Figure 8B-C**). We next asked whether VPS35 variants affect the subcellular localization of TBC1D5 in HEK-293T cells. Using immunofluorescent labeling and confocal microscopy in HEK-293T cells overexpressing V5-tagged VPS35 variants, we evaluated the degree of co-localization between endogenous TBC1D5 and VPS35. We find that TBC1D5 co-localizes equivalently with WT and D620N VPS35, suggesting that the D620N mutation does not alter the distribution of TBC1D5 within the cell (**Figure 8D-E**). Our data suggests that decreased binding of TBC1D5 to D620N VPS35 is not due to the redistribution of TBC1D5 to other cellular compartments devoid of VPS35.

**Figure 8.**
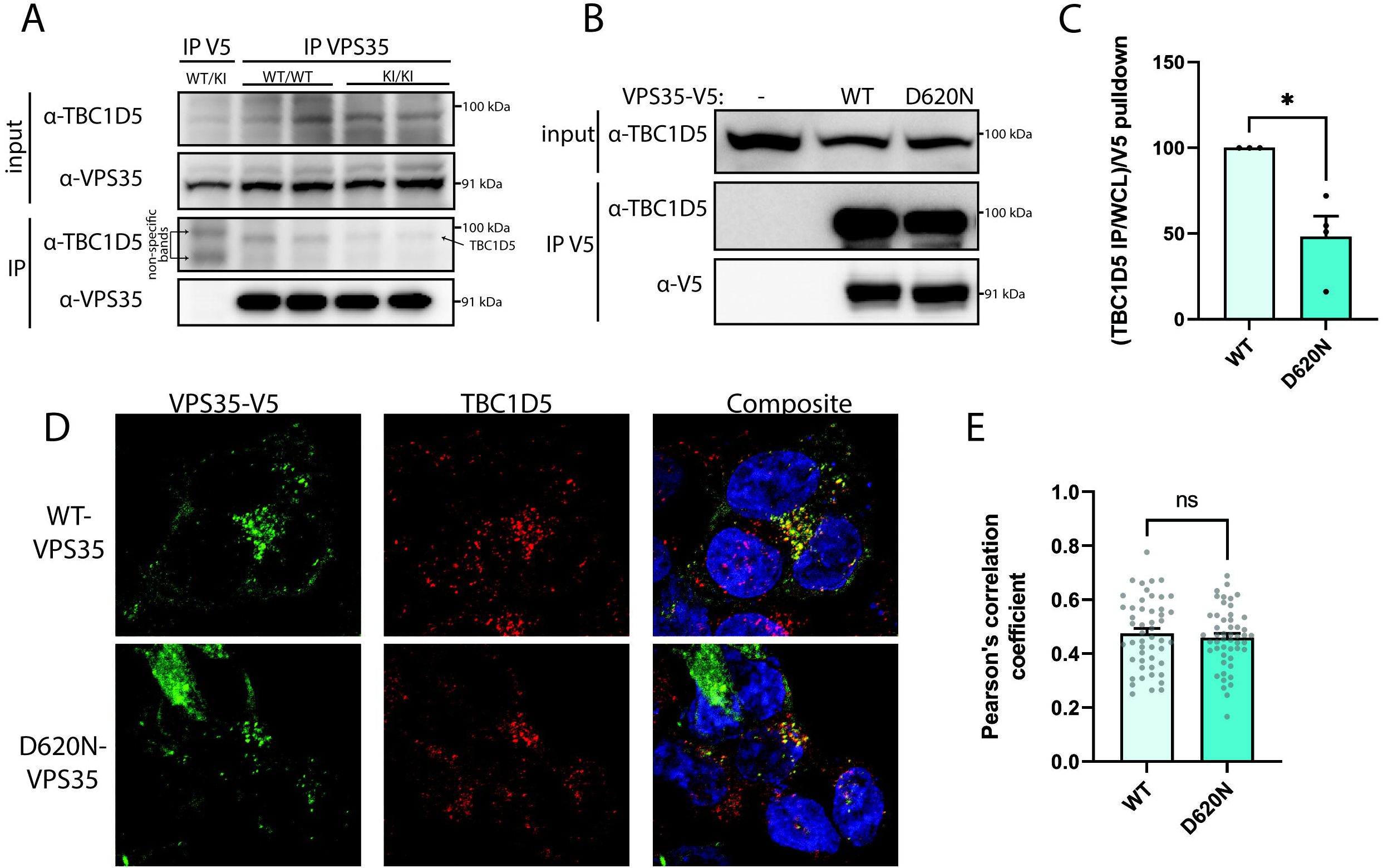
Reduced interaction of TBC1D5 with D620N VPS35. (**A**) Soluble striatal extracts from 3-4 month-old WT and *D620N VPS35* KI mice were subjected to IP with anti-VPS35 antibody (or isotype-matched control anti-V5 IgG) followed by Western blot analysis. Input and IP fractions were probed with anti-VPS35 and anti-TBC1D5 antibodies confirming a TBC1D5 binding deficit in *D620N VPS35* KI brain. Arrows indicate the position of TBC1D5 in the IP VPS35 samples and two non-specific bands in the control IP V5 sample. (**B**) HEK-293T cells expressing V5-tagged VPS35 variants (WT or D620N) were subjected to IP with anti-V5 antibody followed by Western blot analysis. Inputs and IPs were probed with anti-TBC1D5 or anti-V5 antibodies. (**C**) Graph indicates densitometric analysis of TBC1D5 levels in V5 IP, normalized to endogenous levels of TBC1D5 in the input, and expressed as a percent of the WT VPS35 IP condition (mean ± SEM, *n* = 3-4 experiments). (**D**) Representative confocal immunofluorescent images of HEK-293T cells co-labeled for endogenous TBC1D5 (red) and V5-tagged VPS35 variants (green). (**E**) Graph indicates Pearson’s correlation coefficients for TBC1D5 with WT or D620N VPS35 (mean ± SEM, *n* = 50 cells/condition). Data were analyzed by unpaired, two-tailed Student’s *t*-test (\**P*<0.05).

### Impact of the D620N mutation on the interaction of VPS35 with Rab7a

TBC1D5 is a known interacting protein of the retromer and has been shown to function in the endolysosomal system as a GTPase-activating protein (GAP) for Rab7a, a protein responsible for the endosomal recruitment of the retromer (40,41). When activated, GTP-bound Rab7a helps to recruit and bind the retromer to the endosomal membrane (7). TBC1D5 acts to inactive Rab7a, through promoting the hydrolysis of GTP to GDP, thus decreasing the interaction of the retromer with Rab7a at the endosomal membrane (16). Since Rab7a is a direct effector of TBC1D5, we hypothesized that diminished binding of TBC1D5 to D620N VPS35 may also impact Rab7a binding to VPS35. In addition to evaluating the binding capacity of WT and D620N VPS35 to WT Rab7a, we also tested two functional variants of Rab7a. The T22N variant acts as a dominant-negative mutation by reducing the affinity of Rab7a for GTP, effectively rendering Rab7a inactive. The Q67L variant locks Rab7a in its GTP-bound state, making it constitutively active (42).

HEK-293T cells co-expressing V5-tagged VPS35 variants (WT or D620N) and GFP-tagged Rab7a variants (WT, T22N, or Q67L) were subjected to IP with anti-GFP antibody, followed by Western blot analysis with anti-V5 or anti-GFP antibodies. We demonstrate that VPS35 binding to both WT and T22N Rab7a are not affected by the D620N mutation (**Figure 9A-B**). However, WT and D620N VPS35 bind less efficiently to inactive T22N Rab7a compared to WT Rab7a, consistent with prior studies indicating that active Rab7a is responsible for binding to and recruiting VPS35 to the endosomal membrane (**Figure 9A-B**). Notably, we also find that D620N VPS35 significantly increases the binding affinity for constitutively-active Q67L Rab7a compared to WT VPS35 (**Figure 9A-B**). This finding could be consistent with TBC1D5 binding deficits in D620N VPS35 since the presence of the constitutively-active form of Rab7a could recapitulate and compound the proposed effects of TBC1D5 deficiency on Rab7a localization to the membrane upon activation. We also wanted to determine if the steady-state levels of Rab7a are affected by increasing levels of WT or D620N VPS35. HEK-293T cells expressing a constant amount of GFP-tagged Rab7a and increasing amounts of WT or D620N VPS35 were subjected to Western blot analysis. We find a dose-dependent yet equivalent effect of WT or D620N VPS35 on increasing Rab7a levels, suggesting that VPS35 variants in general increase the protein stability of Rab7a (**Figure 9C-D**).

**Figure 9.**
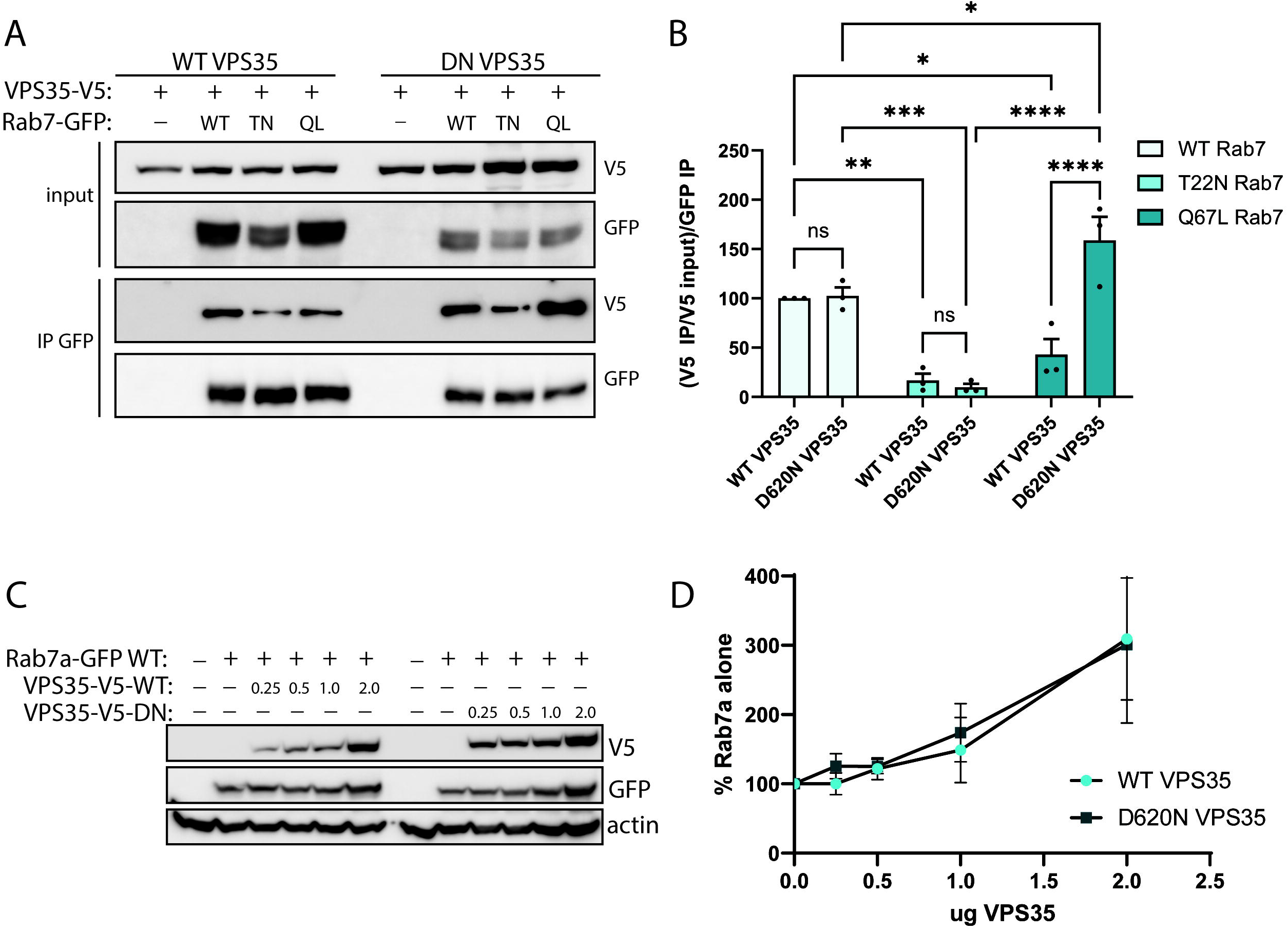
Impact of D620N VPS35 on Rab7 interaction and steady-state levels. (**A**) HEK-293T cells co-expressing V5-tagged VPS35 (WT or D620N) and GFP-tagged Rab7 (WT, T22N, Q67L) were subjected to IP with anti-GFP antibody followed by Western blot analysis. Input and IP fractions were probed with anti-GFP and anti-V5 antibodies. D620N VPS35 displays an increased the interaction with constitutively-active Q67L Rab7. (**B**) Graph indicates densitometric analysis of V5-VPS35 levels in GFP-Rab7 IP, normalized to levels of V5-VPS35 in the input, and expressed as a percentage of WT Rab7 (mean ± SEM, *n* = 3 experiments). Data were analyzed by two-way ANOVA with Tukey’s and Šidák’s multiple comparisons tests (\**P*<0.05, \*\**P*<0.01, \*\*\**P*<0.001, or \*\*\*\**P*<0.0001 as indicated). (**C**) Equivalent Triton-soluble fractions from HEK-293T cells co-expressing GFP-tagged Rab7 and increasing amounts (0.25 – 2 μg plasmid DNA) of V5-tagged VPS35 (WT or D620N) were probed with anti-V5, anti-GFP or anti-actin antibodies to monitor Rab7 steady-state levels. (**D**) Graph indicates densitometric quantitation of Rab7 levels normalized to actin, and expressed as a percent of Rab7 levels alone (mean ± SEM, *n* = 3 experiments).

### Global proteomic analysis of striatal tissue from *WT* and *D620N VPS35* KI mice

We next sought to determine if the D620N VPS35 mutation affects the proteome at a global level in the mouse striatum, potentially indicative of altered retromer cargo sorting and degradation. As such, we rapidly isolated and flash froze the striatum from two WT and two homozygous *D620N VPS35* KI mice. One striatal hemisphere from each animal was subjected to proteome-wide analysis by LC-MS/MS. Tissue was first homogenized and cleared by centrifugation, and 10 μg of protein was resolved by SDS-PAGE and the mobility region of the gel was excised and digested followed by LC-MS/MS analysis. Data were searched using MaxQuant to determine peptide intensity values (43). Statistical analysis reveals a high degree of correlation between all four samples (R > 0.98), suggesting that the D620N VPS35 mutation has a limited impact on the mouse striatal proteome (**Figure 10A**). Similar to our brain co-IP assays, the total proteome of WT and *D620N VPS35* KI striatal tissues are remarkably similar (**Figure 10B**). We successfully identify over 1900 proteins, and 1869 of these proteins are commonly identified between WT and *D620N VPS35* KI mice (**Figure 10B**). We identify a number of known retromer-associated proteins, including cargo-selective complex subunits (VPS35, VPS26A, VPS26B, VPS29), sorting nexins (SNX1, 2, 3, 5, 6, 27), WASH complex subunits (strumpellin), Rab7a, and retromer cargo (GLUT1, GluR1, GluR2), yet none of these proteins are significantly altered in *D620N VPS35* KI striatum (**Figure 10C**). Several proteins are completely absent from either the WT or D620N samples, and due to the low number of proteins that show a significant fold change (**Figure 10C**), these unique proteins are likely the best candidates for further validation.

**Figure 10.**
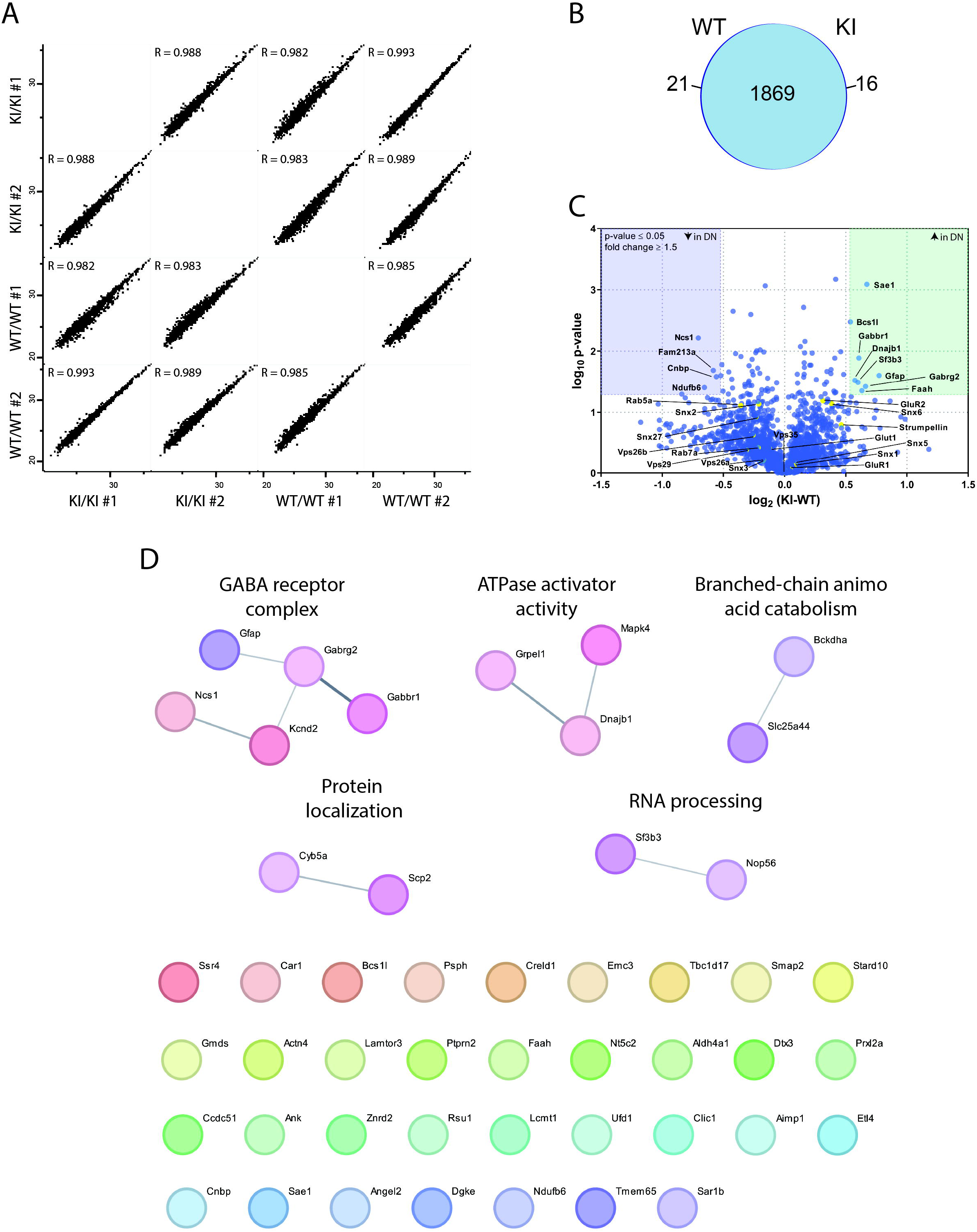
Global proteomic analysis of striatum from WT and *D620N VPS35* KI mice. (**A**) Correlation analysis comparing the log2-transformed intensity values of proteins identified in each condition/replicate compared with all other conditions. All samples were highly correlated (R > 0.98). (**B**) Proportional Venn diagram indicating the strong overlap in global proteins identified in striatum between WT and homozygous *D620N VPS35* KI. (**C**) Volcano plot of LC-MS/MS total proteome data plotted as the log_2_ fold change of WT vs *D620N VPS35* KI striatum against the –log_10_ p-value. Significance was considered for p-values ≤ 0.05 and fold changes greater than 1.5 (*n* = 2 mice/group). (**D**) STRING analysis using MCL clustering (inflation parameter = 3) of a subset of 49 proteins (21 proteins found only in WT VPS35, 16 proteins found only in D620N VPS35, and 12 proteins with significant fold changes) identifies five clusters representing GABA receptor complex, ATPase activator activity, branched-chain amino acid catabolism, protein localization, and RNA processing.

We identify 21 proteins that are completely absent from *D620N VPS35* KI striatum (**Table 6**). Of particular interest are TBC1D17, LAMTOR3 and SMAP2, given their suggested roles within the endolysosomal system (44–46). We also used ORA to compare a subset of 49 proteins (21 proteins found only in WT VPS35, 16 proteins found only in D620N VPS35, and 12 proteins with significant fold changes) against the 1906 proteins identified in total, and no pathways (GO or KEGG) are significantly enriched, further suggesting that the D620N VPS35 mutation does not significantly impact the global striatal proteome. Using STRING analysis with MCL clustering of the subset of 49 proteins, only 14 proteins could be connected to a significant cluster (**Figure 10D**). Clusters include the GABA receptor complex, ATPase activator activity, branched-chain amino acid catabolism, protein localization, and RNA processing. It remains to be determined how VPS35 affects the levels of these proteins or how they might be relevant to mutant VPS35-induced neurodegeneration. Future studies will seek to explore the functional interactions of these proteins with VPS35 and determine their roles in *VPS35*-linked PD.

## Discussion

The studies described here aimed to determine the mechanism(s) through which the PD-linked D620N mutation in VPS35 exerts its pathogenic effects, specifically by focusing on determining key protein interactors that are disrupted by this mutation. It remains unclear how the *D620N VPS35* mutation causes neurodegeneration but several pathways have been implicated, including autophagy/lysosomal disruptions, mitochondrial deficits, and neurotransmitter dysfunction (8,10,11). None of these pathways have been proven to directly cause neurodegeneration, leaving the molecular mechanism of D620N VPS35-induced neurodegeneration unknown. With this in mind, we adopted an unbiased approach to identify differential protein interactors of VPS35 variants in cell-based and rodent models, which identified TBC1D5 as a differential interacting protein of D620N VPS35.

First, we used TAP to identify native protein complexes that interact with VPS35. TAP was an attractive method to study protein-protein interactions since it does not require the use of detergents which allows complexes to stay in their native conformation (12). In principle, due to its gentle nature, TAP should recover interactions that are labile and sensitive to the use of detergents. Additionally, using two consecutive affinity purification steps allows for increased sensitivity and specificity (12,13). This approach has never been used to identify protein complexes that interact with VPS35 in their native state. Interestingly, while able to identify VPS35 and subunits of the retromer cargo-selective complex, our first TAP experiments recovered very few proteins, even after several optimization steps, including changing buffers from Tris-based to HEPES-based, suggesting that VPS35 interactions may be too weak to withstand two consecutive affinity purification steps.

To overcome the difficulties associated with TAP in identifying interacting proteins of VPS35, we optimized the addition of a reversible protein crosslinker to our protocol. We used the reversible and cell permeable crosslinker DSP to crosslink interacting proteins of VPS35 prior to cell lysis to preserve native interactions. Additionally, we anticipated that DSP treatment would preserve weak interactions with VPS35. After optimizing the use of DSP, we were able to substantially increase the number of proteins recovered using TAP, however, we were unable to identify retromer cargo or accessory proteins, suggesting that many of these interacting proteins may be too weak to withstand two consecutive affinity purification steps, even with the addition of a crosslinking reagent. Additionally, we have considered that the TAP-tag itself may interfere with protein binding to VPS35. Since we were able to identify VPS26 and VPS29 in our TAP experiments, we consider such interference unlikely, however, it cannot be excluded that the C-terminal TAP tag may interfere with multimerization of the retromer complex, as described in a recent study demonstrating that VPS35 forms homodimers to encourage flexibility of the retromer during endosomal tubule formation (15). The use of DSP as a crosslinking agent also comes with the caveat that it only reacts with exposed lysine residues at fixed distance within the interface of VPS35 and its interacting proteins, possibly excluding many proteins from being covalently modified.

Next, we used a traditional co-IP approach in HEK-293T cells overexpressing V5-tagged VPS35 variants treated with DSP to determine if we could improve the recovery of interacting proteins in a cell-based model. Indeed, we were able to greatly increase the number of proteins identified by LC-MS/MS analysis in addition to recovering several known interacting proteins of VPS35 outside of the cargo-selective complex, including multiple SNX proteins, the WASH complex subunit strumpellin, and Rab proteins. While we demonstrate that the VPS35 protein interactome in this model is remarkably similar between WT and D620N VPS35 proteins, we nonetheless uncovered several proteins with reduced binding to D620N VPS35 involved in metabolic processes that have not previously been associated with retromer function. Future studies should determine if dysfunction of these pathways contribute to the pathogenic effects induced by D620N VPS35 and the mechanism(s) by which this might occur.

While our cell-based studies provide important insight into the basic biology of the retromer and potential pathways impacted by the D620N VPS35 mutation, it was important to evaluate interactions in brain tissue of disease-relevant animal models. As such, we bilaterally injected the substantia nigra of adult rats with AAV2/6 vectors expressing V5-tagged human VPS35 (WT or D620N) and control virus (MCS) in opposite hemispheres. At 4 weeks post-injection, soluble hemi-brain extracts were used for co-IP and LC-MS/MS analysis, enabling us to explore the impact of the D620N VPS35 mutation selectively within the nigrostriatal dopaminergic pathway. Additionally, our study provided the first characterization of the brain-specific protein interactome of VPS35 and the retromer. As with our cell-based models, WT and D620N VPS35 protein interactomes shared a high degree of similarity in this rodent model, further supporting our assertion that D620N VPS35 remains largely functional (14,22–24). Interestingly, we identified 67 binding proteins that were either absent from or depleted with D620N VPS35, representing both previously described and novel pathways related to the retromer.

While our overexpression-based rat model yielded valuable insight into the biology of the retromer in the brain and the impact of the D620N VPS35 mutation on protein-protein interactions in a disease-relevant tissue, it is based on non-physiological levels of VPS35 expression that does not adequately recapitulate human genetics. While these experiments were being performed, a *D620N VPS35* KI mouse model was developed by our group (38), allowing us to perform interactome and global proteomic studies in a mouse model with endogenous expression of VPS35, making this model the most physiologically-relevant to disease. Through LC-MS/MS analysis, we show that the VPS35 interactome in WT and D620N KI whole brain and striatum are strikingly similar, consistent with data from our overexpression models. Despite the similarity, we identify several proteins that are disrupted by the D620N VPS35 mutation and that require further validation. We demonstrate that TBC1D5 exhibits decreased binding to D620N VPS35 in KI mice compared to WT, in both whole brain and striatal co-IP fractions. We were able to confirm this finding in HEK-293T cells overexpressing human VPS35 variants. We also find that the D620N mutation does not alter the co-localization of VPS35 and TBC1D5 in HEK-293T cells, suggesting that D620N VPS35 does not simply cause a redistribution of TBC1D5 to subcellular localizations devoid of VPS35.

TBC1D5 is a known interacting protein of the retromer and has been shown to function in the endolysosomal system as a GTPase-activating protein (GAP) for Rab7a, a known regulator of retromer localization to endosomal membranes (7,40,41). TBC1D5 interacts with the retromer through a strong association with VPS29 and a weaker association with VPS35. We observe decreased binding of VPS29 to D620N VPS35 in our brain KI co-IP assays, which could explain downstream TBC1D5 binding deficits. We hypothesized that TBC1D5 binding deficits with D620N VPS35 would also affect Rab7a binding to VPS35. While we do not find a difference in Rab7a binding to WT or D620N VPS35 in HEK-293T cells, we do find an increased binding affinity of D620N VPS35 for constitutively-active Q67L Rab7a compared to WT VPS35. This finding could suggest that Q67L Rab7a phenocopies the effects of reduced TBC1D5 binding in that both conditions lock active Rab7a on late endosomal membranes (47). This could be consistent with a prior study suggesting the association of TBC1D5 with the retromer increases its GAP activity towards Rab7a, which is essential for the sorting of CI-MPR (41). This could also align with studies suggesting that the D620N mutation alters the recycling of CI-MPR (22,48,49).

On the other hand, TBC1D5 inactivation of Rab7a has been suggested to decrease the localization of retromer to endosomal membranes (16). Our data raise the possibility that overexpression of TBC1D5 may be able to restore binding deficits induced by the D620N VPS35 mutation. However, contradictory to this idea is a study suggesting that loss of TBC1D5 can restore the WASH complex binding deficits observed for D620N VPS35 (16). It was shown that TBC1D5 together with the retromer serves to regulate the localization and activity of Rab7a (50). Under normal conditions, Rab7a is widely localized throughout the cell, decorating endosomes, lysosomes, mitochondria, endoplasmic reticulum and the *trans*-Golgi network. Loss of TBC1D5 or the retromer induced a distinct endolysosomal accumulation of GTP-bound, active Rab7a (50). Without the control of TBC1D5 and the retromer, Rab7a becomes hyperactive and immobile on endolysosomal membranes, essentially decreasing the pool of inactive Rab7a within the cell. This impact on Rab7a may lead to defects in ATG9A sorting and parkin-dependent mitophagy (50). Several questions remain regarding TBC1D5 and the retromer and the role this interaction plays in neurodegeneration. Specifically, it remains unclear exactly why TBC1D5 interacts so strongly with the retromer when its interaction serves to dissociate the retromer from the endosome via Rab7a inactivation. Perhaps the interaction between TBC1D5 and the retromer is more complex than originally thought and TBC1D5 acts as a regulator of retromer function through another mechanism or GAP effector. Future experiments will aim to elucidate the role of VPS29, TBC1D5 and Rab7a in mutant VPS35-induced neurodegeneration.

Another important observation from our brain co-IP assays was the inability to recover any components of the WASH complex, which is known to exhibit a decreased interaction with D620N VPS35 via binding of the Fam21 subunit (14,22). Since this interaction has yet to be validated in the brain, we hypothesized that co-IP experiments from the brain of D620N VPS35 KI mice would be able to replicate WASH complex binding deficits. Surprisingly, we were unable to recover any WASH complex components in our brain co-IP assays, even though a previous study from our group indicates abundant levels of both WASH1 and Fam21 subunits in soluble mouse whole brain extracts (38). We were also able to recover WASH complex components in our striatum global proteome study, further suggesting that WASH complex proteins are detectable in the brain. As we were unable to recover this interaction, it is possible that the WASH complex is not a significant binding partner of VPS35 in the brain, raising the interesting possibility that the reported WASH complex binding deficit is not relevant in the context of brain pathology in PD. Further work to validate our brain-specific co-IP findings will be necessary to understand the pathogenic effects of the D620N VPS35 mutation.

While WT and D620N VPS35 protein interactomes demonstrated remarkable similarity within each experimental paradigm, marked differences were noted between experimental paradigms. Of 870 total interacting proteins that met our significance criteria in at least one of our brain-specific co-IP assays, only 49 proteins were commonly identified in all three paradigms, suggesting that the VPS35 interactome is regulated in a tissue– and cell type-specific manner, and likely further regulated by endogenous versus overexpression levels of VPS35. However, overrepresentation analysis of the 49 common interacting proteins provides insight into the conserved functions of the retromer in cellular homeostasis in brain tissue, regardless of specific brain region, cell type or expression level. These data are supported by a previous study from our group demonstrating early post-natal lethality and neurodegeneration in a pan-neuronal conditional *VPS35* knockout mouse model (51). Further studies will be needed to dissect the cell-specific roles of VPS35 in different brain regions and cell types. Importantly, we provide the first characterization of the endogenous VPS35 protein interactome in the brain, providing insight into the role of the retromer in PD-relevant tissue.

Finally, we evaluated the striatum of WT and KI mice for global proteomic changes induced by the VPS35 D620N mutation. As with the brain IP interactome screen, we reveal limited differences between the striatal proteomes of WT and D620N VPS35 KI mice, suggesting that the D620N mutation does not disrupt protein levels on a wide scale. We were able to identify several proteins whose levels were altered by the D620N VPS35 mutation, however, further validation will be required to determine how these proteins might contribute to mutant VPS35-induced neurodegeneration. Of these proteins, further validation of TBC1D17, LAMTOR3 and SMAP2 are warranted based upon their absence from the D620N VPS35 KI striatum and their known roles in the endolysosomal system (44–46). It is not yet known how *VPS35* mutations affect the levels or function of these proteins, whether they are also altered in other brain regions beyond the striatum, or if they are relevant to mutant VPS35-induced neurodegeneration.

Taken together, our data provides a comprehensive evaluation of proteomic changes induced by the D620N mutation in brains of *VPS35* KI mice. We identified TBC1D5 as a promising differential interacting protein of VPS35. Future experiments will focus on understanding how TBC1D5 contributes to mutant *VPS35*-linked neurodegeneration with the aim of elucidating how the *D620N VPS35* mutation drives neurodegeneration in PD.

## Materials and Methods

### Animals

*D620N VPS35* KI mice (stock 021807) were originally obtained from The Jackson Laboratory, crossed with CMV-Cre transgenic mice (strain 006054) to remove a floxed “mini-cDNA” containing wild-type *VPS35* exons 15-17, and the resulting germline KI have been described previously (38). Adult female Sprague Dawley rats (175-200 grams) were obtained from Charles River. Animals were maintained in a pathogen-free barrier facility and provided with food and water *ad libitum* and exposed to a 12 h light/dark cycle. Animals were treated in strict accordance with the NIH Guide for the Care and Use of Laboratory Animals. All studies using mice and rats were reviewed and approved by the Institutional Animal Care and Use Committee (IACUC) of Van Andel Institute under protocols AUP# 23-07-017 and 23-11-030.

### Expression plasmids and antibodies

A mammalian expression plasmid (pLenti6-V5/DEST) containing full-length human VPS35 with a C-terminal V5 tag was obtained from Addgene (#21691). D620N VPS35 was generated as previously described (24). TAP-tagged VPS35-WT was generated using restriction cloning to ligate a pENTR/D-TOPO-VPS35 vector with a 3’ TAP tag fragment (∼218 nucleotides) derived from a pCTAP vector (Agilent). TAP-tagged VPS35-D620N was created by site-directed mutagenesis (Agilent QuickChange II XL kit) and sequenced to confirm integrity. The resulting pENTR-VPS35-CTAP entry vectors were subjected to Gateway-based recombination into a pcDNA3.1/V5-DEST destination vector. Rab7a plasmids containing full-length human Rab7a with a N-terminal EGFP and 2xHA tag were obtained from Addgene (WT, #28047; T22N, #28048; Q67L, #28049). The following primary antibodies were used: mouse anti-VPS35 (ab57632, Abcam), mouse anti-VPS35 (SMC-602, stressmarq), mouse anti-V5 (Thermo Fisher), mouse anti-V5-HRP (R96125, Invitrogen), mouse anti-GFP (clone 7.1 and 13.1, Roche), rabbit anti-VPS26 (ab23892, Abcam), goat anti-VPS29 (ab10160, Abcam), rabbit anti-WASH1 (SAB4200372, Sigma-Aldrich), rabbit anti-Rab7 (ab137029, Abcam), mouse anti-actin (MAB1501, Millipore), mouse anti-β-tubulin (Sigma, T5201), rabbit anti-TBC1D5 (ab203896, Abcam) and mouse anti-TBC1D5 (sc-376296, Santa Cruz Biotechnology). Secondary HRP-conjugated antibodies used for Western blotting were: goat anti-mouse IgG, light chain-specific (Jackson Immunoresearch), mouse anti-goat IgG, light chain-specific (Jackson Immunoresearch) and mouse anti-rabbit IgG, light chain-specific (Jackson Immunoresearch). For confocal fluorescence analysis, the following secondary antibodies were used: AlexaFluor-488 or –647 goat anti-mouse IgG (H + L) and AlexaFluor-594 goat anti-rat IgG (H + L) (Thermo Fisher).

### Cell culture and transient transfection

Human HEK-293T or SH-SY5Y cells were maintained at 37°C with 5% CO_2_ in Dulbecco’s modified Eagle’s media (DMEM) (Gibco) supplemented with 10% fetal bovine serum and penicillin/streptomycin. Transient transfection was performed with plasmid DNAs using XtremeGene HP DNA Transfection reagent (Roche) according to the manufacturer’s instructions.

### Tandem affinity purification

The commercially available InterPlay Mammalian TAP system (Agilent Technologies) was used according to manufacturer’s instructions, however, buffers were made inhouse using HEPES to identify VPS35-interacting proteins. Briefly, HEK-293T cells were transiently transfected with TAP-tagged VPS35 (WT or D620N) or empty vector. If treated with DSP, cells were treated with 0.3 mM DSP diluted in PBS for 30 min at room temp. The cross-linker reaction was quenched at RT for 10 min with 1 M glycine. Cells were resuspended in TAP lysis buffer (0.1% NP-40, 10% glycerol, 50 mM HEPES-KOH, 100 mM KCl, 1X Complete Mini protease inhibitor cocktail [Roche]) and lysed via successive rounds of freeze-thaw cycles. To each lysate, 2 mM EDTA was added before incubation with streptavidin resin at 4°C overnight. The resin was rinsed twice with streptavidin binding buffer (TAP lysis buffer + 2 mM EDTA) followed by a 4-hour incubation with the biotin containing streptavidin elution buffer (streptavidin binding buffer + 4 mM biotin). Next, the eluate containing the streptavidin-purified proteins was incubated overnight at 4°C with a calmodulin resin in calmodulin binding buffer (TAP lysis buffer supplemented with 1 mM magnesium acetate, 1 mM imidazole, and 2 mM CaCl_2_). Finally, the resin was rinsed twice in calmodulin binding buffer before proteins were eluted from the calmodulin resin by first incubating the resin with 2X sample buffer supplemented with 50 mM DTT for 10 min at room temp, followed by a 5-min incubation at 95°C. The final elution was sent for LC-MS/MS analysis as described below.

### Silver staining of protein gels

To validate TAP eluates, the SilverQuest Silver Staining Kit (Invitrogen) was used according to manufacturer guidelines. Following electrophoresis, SDS-PAGE gels with TAP eluates were briefly rinsed with ddH_2_O before fixing overnight at room temperature in fixation solution (40% ethanol, 10% acetic acid). The gel was then washed in 30% ethanol for 20 min, followed by an incubation with Sensitizing Solution (30% ethanol, 10% Sensitizer) for 20 minutes. Following a 20-min ddH_2_O wash, the gel was then incubated with Staining Solution (1% Stainer) for 30 min. After a brief 30 sec wash in ddH_2_O, the gel was then incubated in Developing Solution (10 ml Developer, 1 drop Developer Enhancer, plus 90 ml water) until the desired band intensity was reached (approximately 10 min). Once the desired band intensity was reached, 10 ml of Stopper Solution was added directly to the Developing Solution and incubated for 10 min. Finally, the gel was washed for 10 min in ddH_2_O before imaging on a ChemiDoc Imaging System (Bio-Rad).

### Co-immunoprecipitation and Western blotting

For brain-specific co-IP assays, fresh brain tissue (hemi-brains or striatum) from 3-4 month old mice were homogenized in Tris– or HEPES-based lysis buffer (Tris-buffer: 50 mM Tris-HCl pH 7.5, 150 mM NaCl, 1% Triton X100, 5% glycerol, 1 mM EDTA and 1X Complete protease inhibitor cocktail [Roche]; HEPES-buffer: 20 mM HEPES-KOH, pH 7.2, 50 mM potassium acetate, 200 mM sorbitol, 2 mM EDTA, 0.1% Triton X-100, 1X Complete Mini protease inhibitor cocktail [Roche]). Homogenized brain tissue was centrifuged at 100,000 x *g* for 30 min at 4°C, after which equivalent soluble fractions were incubated overnight with Protein G-Dynabeads (Thermo Fisher) that had been pre-incubated with anti-VPS35 (5 ug) antibody (or anti-V5 [5 ug] antibody as an isotype-matched antibody control). After an overnight incubation, Dynabeads were washed twice with lysis buffer and twice with lysis buffer without Triton X-100. IPs were eluted by boiling at 95°C for 5 min in 50 µl 2X Laemmli sample buffer (Biorad). Inputs and IPs were subjected to SDS-PAGE, transferred to nitrocellulose membranes, followed by Western blot analysis and imaging on a FujiFilm LAS-4000 Luminescent Image Analysis system.

For cell-based co-IP assays, HEK-293T cells were transiently transfected with the desired combination of plasmids in 10 cm dishes. At 48 h post-transfection, cells were harvested in 1 ml of lysis buffer (see above). Cell lysates were allowed to rotate for 1 h at 4°C after which cell lysates were centrifuged at 15,000 rpm for 15 min at 4°C to collect the soluble fraction. Soluble fractions were mixed with Protein G-Dynabeads (Thermo Fisher) that had been pre-incubated with anti-V5 (1 μg) antibody and incubated overnight at 4°C. All co-IP assays were washed as follows: 1x with lysis buffer supplemented with 150 mM NaCl, 2x with lysis buffer, and 2x with lysis buffer without 0.1% Triton X-100. IPs were eluted by boiling at 95°C for 5 min in 50 µl 2X Laemmli sample buffer (Biorad). Inputs and IPs were subjected to SDS-PAGE, transferred to nitrocellulose membranes, followed by Western blot analysis and imaging on a FujiFilm LAS-4000 Luminescent Image Analysis system. Where applicable, densitometry to quantify levels of proteins was performed using Image Studio Lite (LI-COR Biosciences).

### Mass spectrometry and proteomic analysis

Brain co-IP assays were conducted as described above. 10 μl of each IP was used for Western blot analysis and the remaining 40 μl of IP was sent for LC-MS/MS analysis (MS Bioworks, Ann Arbor, MI). Briefly, gel fragments were digested with trypsin and analyzed by nano-LC-MS/MS with a Waters NanoAcquity HPLC system interfaced to a ThermoFisher Q Exactive. Peptides were loaded on a trapping column and eluted over a 75 µm analytical column at 350 nL/min with a 1 h reverse gradient; both columns were packed with Jupiter Proteo resin (Phenomenex). The mass spectrometer was operated in data-dependent mode, with Orbitrap operating at 60,000 FWHM and 17,500 FWHM for MS and MS/MS, respectively. The fifteen most abundant ions were selected for MS/MS. Data were searched using Mascot and parsed into Scaffold software for validation, filtering and to create a non-redundant list per sample. Data were also searched using Maxquant to obtain peptide intensity values. Data were filtered using a 1% protein and peptide FDR and required at least two unique peptides per protein. Data are shown as peptide intensity values.

GO and KEGG pathway analysis was done using the functional annotation tool DAVID (20,21). Bubble plots and Venn diagrams were generated using SRplot (52). Statistical analysis of MS data was performed using the statistical environment R and GraphPad Prism (v.10.2) (53). Functional enrichment analysis was done using the Cytoscape (v.3.10.2) plug-in ClueGo with GO biological process terms (18). Pathways with a p-value of <0.05 and a Kappa score of 0.4 were mapped. Two-sided hypergeometric statistical analysis test with Bonferroni step-down p-value correction was used. Redundant GO terms with >50% overlap were merged and the Leading Group term was determined based on highest significance. Node size corresponds to p-value. When appropriate, STRING diagrams were generated using the open-source website www.string-db.org (54). If applicable, MCL clustering (inflation parameter = 3) was employed.

### Biochemical analysis of tissue

Brains from 3-4 month-old WT or *D620N VPS35* KI mice were regionally dissected and homogenized in lysis buffer (10% w/v) with 50 mM Tris-HCl pH 7.5, 150 mM NaCl, 1% Triton X-100, 5% glycerol, 1 mM EDTA and 1X Complete protease inhibitor cocktail (Roche) as previously described (55). Briefly, the soluble fraction was obtained by centrifugation of homogenized lysates at 4°C for 30 min at 100,000 x *g*. BCA assays (Pierce Biotechnology) were used to determine protein concentration so that equal levels of protein could be loaded and resolved using SDS-PAGE. Western blot analysis was then performed with desired antibodies.

### AAV vectors and stereotactic surgery

Recombinant AAV2/6 viral vectors expressing V5-tagged human VPS35 (WT or D620N) were purified and titered by the University of North Carolina’s Viral Vector Core, as previously described (56). Stereotactic injections in adult female Sprague Dawley rats (175-200 grams) were performed as previously described (57). Briefly, bilateral injections were performed using the following coordinates relative to bregma: anterior-posterior: –5.2 mm, mediolateral: –2 and +2 mm, and dorsoventral (DV): –7.8 mm, to deliver AAV2/6-VPS35 (WT or D620N) and AAV2/6-MCS to opposite hemispheres of the substantia nigra pars compacta. Each rat received ∼5 x 10^9^ viral genomes (vg) of AAV2/6 vector in a volume of 3 µl at a flow rate of 0.2 µl/minute, per hemisphere. Rats were sacrificed at 4 weeks post-injection and hemi-brain tissue was subjected to co-IP with anti-V5 antibody (Thermo Fisher), Western blotting, and mass spectrometry analysis.

### Immunocytochemistry

Transiently transfected HEK-293T cells seeded on glass coverslips in 35 mm dishes were fixed in 4% paraformaldehyde (PFA) and processed for immunocytochemistry analysis as previously described (58). Briefly, following fixation, coverslips were incubated with primary antibodies overnight at 4°C, followed by incubation with AlexaFluor-conjugated secondary antibodies at room temperature for 2 h. Coverslips were mounted onto glass slides using Prolong Diamond Antifade Mountant with DAPI (Thermo Fisher). Fluorescent images were acquired in x, y, and z planes using a Nikon A1plus-RSi Laser Scanning Confocal microscope (Nikon Instruments) equipped with a 60x oil objective. Images were subjected to deconvolution using Huygens Professional (Scientific Volume Imaging). Co-localization coefficients of maximum intensity projection images were calculated using Nikon Elements (Nikon Instruments).

### Statistical analysis

Statistical analysis of protein densitometry was performed using GraphPad Prism 10 (RRID:SCR_002798) by unpaired, two-tailed Student’s *t*-test or by two-way ANOVA with Tukey’s and Šidák’s multiple comparisons tests. Graphs were generated using GraphPad Prism 10 and all data displayed as mean ± SEM. Two-tailed Student’s *t*-test of protein intensity was performed in Microsoft Excel and Volcano plots were generated using GraphPad Prism 10. Correlation analysis was performed using Pearson’s correlation in R (v.4.5.1), and results were visualized using ggplot2 with linear regression fits. DAVID was used to determine the top GO and KEGG terms, based on Bonferroni-corrected p-value and fold enrichment and displayed as bubble plots generated using SRplot (20,21,52). Functional enrichment analysis using the Cytoscape plug-in ClueGo with GO terms was performed with mapping based on pathways with a p-value of <0.05 and a Kappa score of 0.4 (18). Two-sided hypergeometric statistical analysis test with Bonferroni step-down p-value correction was used. Overrepresentation analysis was performed in R using the clusterProfiler package (53). When appropriate, STRING diagrams were generated using the open-source website www.string-db.org (54). If applicable, MCL clustering (inflation parameter = 3) was employed.

## Acknowledgements

The authors thank Dr. Michael Ford (MS Bioworks, Ann Arbor, MI) for his support with mass spectrometry and proteomic analysis. We thank the Van Andel Institute Bioinformatics and Biostatistics Core (RRID:SCR_024762), especially Zach Madaj and Kaitlyn DenHaan, for their assistance with mass spectrometry data analysis. Support for confocal imaging and analysis was provided by Corinne Esquibel of the Van Andel Institute Optical Imaging Core (RRID:SCR_021968). Animal husbandry was provided by the Van Andel Institute Vivarium Core (RRID:SCR_023211). We also thank Dr. Matthew Seaman (Cambridge Institute for Medical Research, University of Cambridge) for his helpful comments regarding buffer components.

## Author contributions

E.T.W. and D.J.M. wrote the main manuscript text and prepared figures. E.T.W., X.C., J.R., S.I. and M.F. designed and conducted the experiments. E.T.W., X.C. and S.I. performed the data analysis. E.T.W. and D.J.M. conceptualized the research. D.J.M. supervised the research.

## Availability of data and materials

All data and protocols generated in this study are included in this manuscript. Additional data and materials are available upon request.

## Declarations

### Ethics approval and consent to participate

All studies using mice and rats were reviewed and approved by the Institutional Animal Care and Use Committee (IACUC) of Van Andel Institute under protocols AUP# 23-07-017 and 23-11-030.

## Consent for publication

All authors have read and approved the final manuscript.

## Supplemental Data

**Figure S1.** STRING clustering of VPS35-interacting proteins altered in rat brain injected with AAV-VPS35. (**A**) STRING analysis using MCL clustering (inflation parameter = 3) of the most abundantly represented pathways from 67 interacting proteins either absent or depleted in the D620N VPS35 condition relative to WT VPS35. Enriched pathways include RHOBTB2 GTPase cycle, the β-catenin destruction complex, mRNA processing, and actin cytoskeleton. (**C**) STRING analysis using MCL clustering (inflation parameter = 3) of the most abundantly represented pathways from 19 proteins either absent or depleted in the WT VPS35 condition relative to D620N VPS35. No significant clustering was identified.

