## Supplementary figures and images for "Parkinson’s disease-linked D620N mutation selectively alters the brain-specific protein interactome of VPS35"

### Figure S1

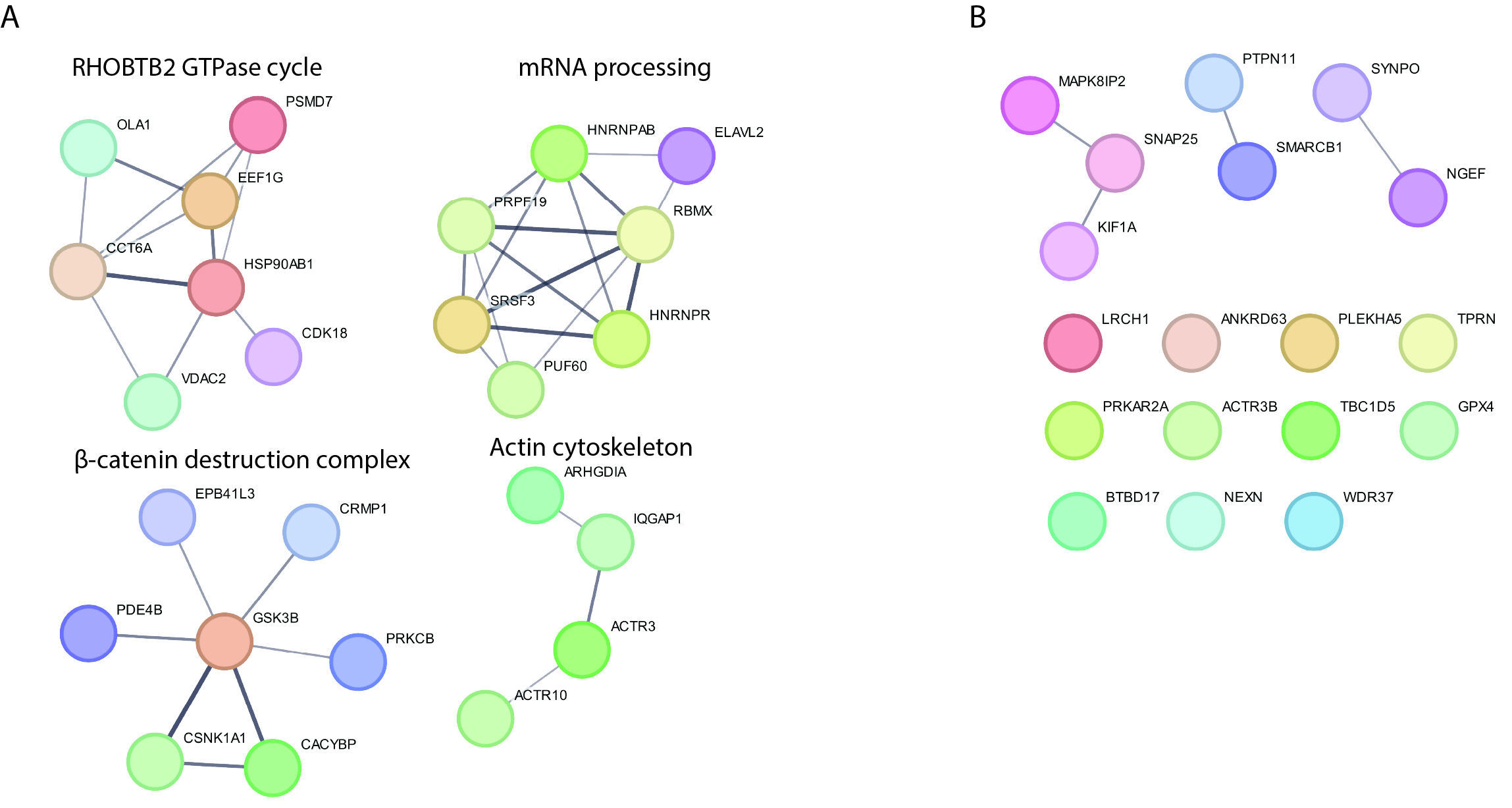
